# Lack of Fe(II) transporters in basal Cyanobacteria complicates iron uptake in ferruginous Archean oceans

**DOI:** 10.1101/2021.11.08.467730

**Authors:** Tristan C. Enzingmüller-Bleyl, Joanne S. Boden, Achim J. Herrmann, Katharina W. Ebel, Patricia Sánchez-Baracaldo, Nicole Frankenberg-Dinkel, Michelle M. Gehringer

## Abstract

Cyanobacteria oxygenated Earth’s atmosphere during the Great Oxygenation Event (GOE) through oxygenic photosynthesis. Their high iron requirement was presumed met by high levels of Fe(II) in the anoxic Archean ocean. **Here we show that** most basal Cyanobacteria cannot synthesize the primary Fe(II) transporter, FeoB. Relaxed molecular clock analyses estimate the arrival of FeoB, as well as the Fe(III) transporters, cFTR1 and FutB, in the Cyanobacteria after the GOE. Furthermore *Pseudanabaena* sp. PCC7367, a basal marine, benthic strain grown under simulated Archean conditions, constitutively expressed *cftr1*, even after the addition of Fe(II). By utilizing gene expression studies under a simulated Archean atmosphere, as well as comparative genomics, phylogenetics and molecular clock analyses, this study identified a need to reappraise iron uptake in ancestral Cyanobacteria, as genetic profiling suggests that scavenging of siderophore bound Fe(III), rather than Fe(II), appears to have been the means of iron acquisition prior to the GOE.

## Introduction

Iron is the fourth most abundant element in the Earth’s crust and is indispensable for life, constituting an essential component in enzymes involved in nitrogen fixation, pigment synthesis, cellular respiration and DNA biosynthesis, to name a few (Sestok *et al*., 2018). It is used more commonly in prokaryotes than any other transition metal (Zerkle *et al*., 2005) and Fe-binding proteins have been preferentially retained in prokaryotic genomes through time, implying it is essential for cellular biochemistry (Dupont *et al*., 2006). Under current aerobic conditions, Fe exists in complex oxides as Fe(III), which are insoluble and kinetically inert. Therefore, low bioavailability of iron in modern oceans has been considered the major factor limiting open ocean primary productivity (Jiang *et al*., 2020a, 2020b). The uptake of iron by prokaryotes has been extensively studied and yet the understanding of the mechanisms, and identification of all the participating receptor components, are still unclear (Fresenborg *et al*., 2020, Qui et al., 2021). Additionally, most studies investigating iron uptake have focused on iron limiting conditions, usually under the oxidizing environment of our present atmosphere, where mechanisms utilizing siderophores, low molecular weight biologic metal chelators that bind free Fe(III) and facilitate its targeted uptake across the prokaryotic cell membranes, dominate (Kranzler *et al*., 2011; Kranzler *et al*., 2014; Årstøl and Hohmann-Marriott 2019; Fresenborg *et al*., 2020).

Early Earth had an anoxic, slightly reducing atmosphere which meant that iron was present as Fe(II) with concentrations approximating 120 µM in the Archean oceans (Canfield, 2005; Catling and Zahnle, 2020). This was significantly altered with the oxygenation of the Earth’s atmosphere during the GOE whereby the redox status of the planet tipped to an oxidizing environment. Cyanobacteria, the only present-day prokaryotes capable of conducting oxygenic photosynthesis, are largely accepted to have generated the copious amounts of oxygen required to oxygenate not only the atmosphere, but also the oceans (Jiang *et al*., 2020a, 2020b; Schopf and Kudryavtsev, 2012). While the GOE is timed at approximately 2,3 – 2,5 Ga (Konhauser *et al*., 2011; Bekker *et al*., 2004; Gumsley *et al*., 2017), signs of large shallow water, phototrophic tidal mats, thought to be ancient Cyanobacteria, appear to have existed at ∼ 3.2 Ga (Heubeck *et al*., 2016; Homann *et al*., 2018) and molecular clocks predict that ancestral cyanobacteria appeared more than a billion years before the GOE (Boden *et al*., 2021; Oliver *et al*., 2021; Cardona, 2019).

Cyanobacteria in general contain a higher metal content than chemoheterotrophic micro-organisms (as summarized in reviews by Fresenborg *et al*., 2020 & Qiu *et al*., 2021). Cyanobacteria can have 25 – 350 times more atoms of iron per cell than *Escherichia coli*, depending on strain, cell type and function (Fresenborg *et al*., 2020). The redox status of Cyanobacteria is tightly coupled to the light cycle, with genes encoding high-affinity metal transporters for iron, manganese and copper following a diurnal expression pattern (Botello-Morte *et al*., 2014; Saha *et al*., 2016). In order for iron to enter the cyanobacterium, it has to cross the outer cell membrane, pass across the periplasmic space and be transported across the inner plasma membrane. An overview of iron specific transporters identified in Cyanobacteria is presented in Figure 1. An extensive summary of iron transporters in Cyanobacteria is reviewed in Fresenborg *et al*., (2020) and Qui *et al*., (2021). Cyanobacterial porins permit the selective passage of compounds through the outer cell membrane, with an iron specific porin recently being identified in *Synechocystis* sp. PCC6803 (Qui *et al*., 2021). Most iron in modern day aquatic systems is bound to organic ligands, siderophores, and crosses the outer membrane via TonB dependent proteins (TBDT) energized by the ExbB/D system on the inner cell membrane (Fig. 1; Fresenborg *et al*., 2020; Qui *et al*., 2021 for reviews). The synthesis of siderophores is not prevalent in the deep rooting basal lineages of Cyanobacteria (Årstøl and Hohmann-Marriott 2019), whereas basic siderophore transporters are commonly identified in a wide range of Cyanobacterial genomes (reviewed by Fresenborg *et al*., 2020 & Qiu *et al*., 2021). To date, Cyanobacterial siderophore synthesis studies have focused on aquatic strains grown under iron depleted conditions, such as those found in the modern open ocean (Årstøl and Hohmann-Marriott 2019)

**Fig. 1.**
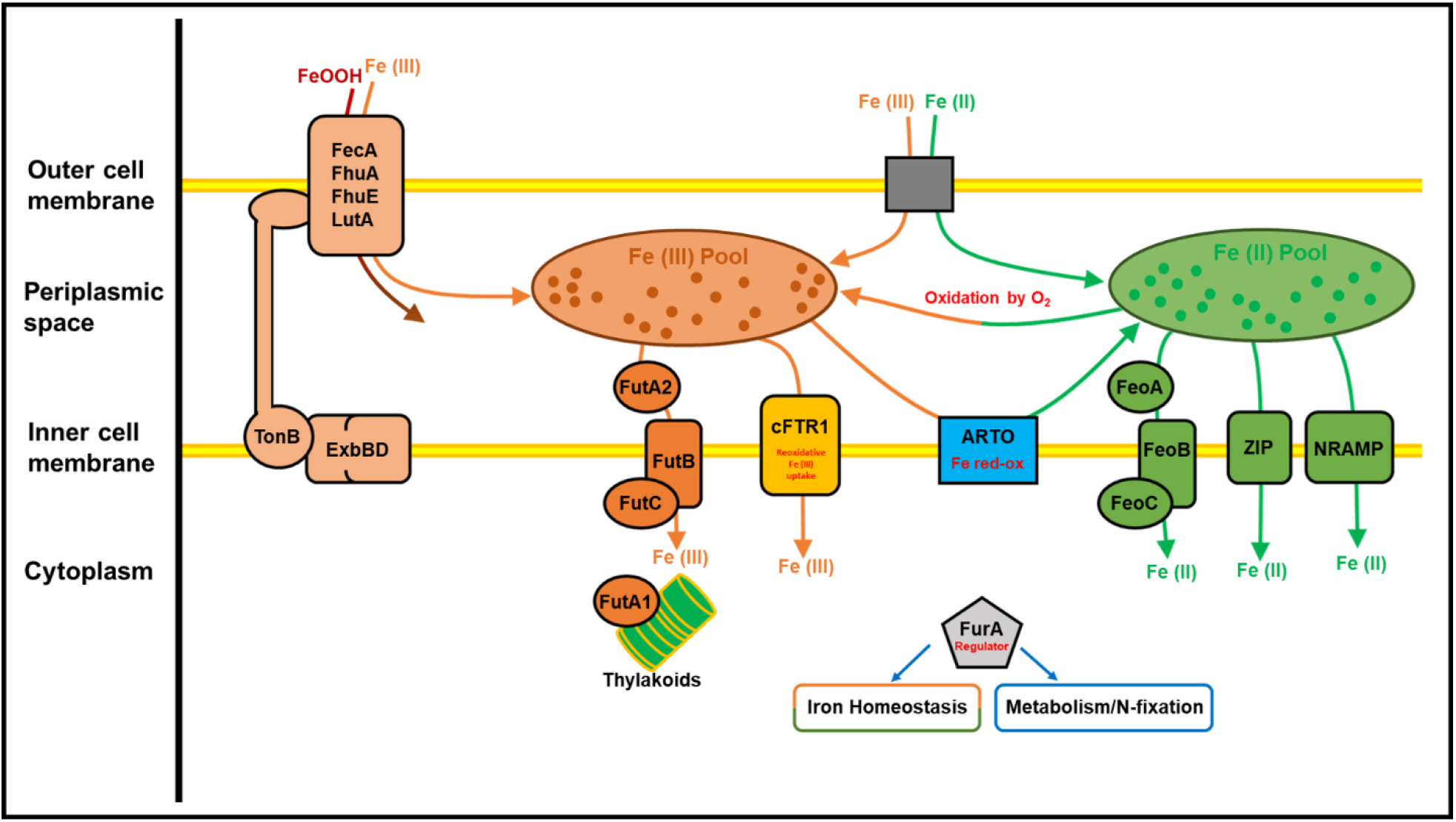
Inorganic iron uptake in Cyanobacteria. Cyanobacteria make use of specific transporters to facilitate inorganic iron uptake: The FutABC system facilitates Fe(III) uptake into the cell (Katoh *et al*., 2001; Brandt *et al*., 2009), while FeoB is the primary transporter for Fe(II) (Katoh *et al*., 2001; Kranzler *et al*., 2014). The zinc-iron permease (ZIP) (Morrissey and Bowler 2012) and the natural resistance associated macrophage protein (NRAMP) homologue (Nevo and Nelson 2006) are thought to take up Fe(II) and other divalent cationic metals. Alternative respiratory oxidases (ARTOs) can mediate the redox state of periplasmic Fe (Hart *et al*., 2005; Berry *et al*., 2002; Katoh *et al*., 2000) and is suggested to play a role in periplasmic iron reduction (Kranzler *et al*., 2011; Kranzler *et al*., 2014). The cytosolic protein, Fur A, functions as a transcriptional repressor (Kaushik *et al*., 2016), regulating iron uptake (González *et al*., 2012; González *et al*., 2013; González *et al*., 2014; González *et al*., 2016). The permease, cFTR1, takes up Fe(III) possibly by re-oxidation of Fe(II) by oxygen (Xu *et al*., 2016). Potentially the TonB – ExbB/D complex co-ordinates with TonB dependent transporters (TBDTs) to take up iron that is bound to organic ligands (Boukhalfa & Crumbliss 2002). ExbB/D was also found to take up inorganic iron directly (Qiu *et al*. 2018).

The current consensus is that cyanobacterial Fe uptake mechanisms emerged from the pre-existing genetic pool of bacterial transporters for reduced metals, namely FeoB, and oxidized iron, cFTR1, TonB and ExbB/D (Qui *et al*., 2021; Fresenborg *et al*., 2020; Jiang *et al*., 2020a). Given the diversity of iron transporters identified in Cyanobacteria, we investigate, in this study, the expression of iron specific transporters under iron replete conditions representing a ferruginous ocean under an anoxic atmosphere. Most investigations into iron transport in Cyanobacteria have focused on the freshwater, unicellular, *feoB* carrying *Synechocystis* sp. PCC6803 (Reviews from Fresenborg *et al*., 2020 and Qiu *et al*., 2021 and references therein). This strain evolved in the completely oxygenated oceans of the Phanerozoic (Sánchez-Baracaldo *et al*., 2015) and therefore offers limited insight into possible processes in the former ferruginous oceans of the Archean. Its genome is also strongly reduced in size compared to modern-day descendants of basal cyanobacterial strains, such as *Pseudanabaena* sp. PCC7367, that emerged 0.95 to 1.25 Ga (Schirrmeister *et al*., 2013; Sánchez-Baracaldo *et al*., 2017; Sánchez-Baracaldo, 2015; Boden *et al*., 2021) and may have retained ancestral metabolisms, including iron uptake mechanisms.

Recently, we found that *Pseudanabaena* sp. PCC7367 was able to survive repeated nocturnal influxes of Fe(II) under anoxic conditions, whereas another deep branching marine strain, *Synechococcus* sp. PCC7336, did not (Herrmann *et al*., 2021). Previous analysis of 72 Cyanobacterial genomes by Kranzler *et al*., (2014; SI Fig. 4), indicated that the genomes of a large number of marine species, including picocyanobacteria and *Pseudanabaena* sp. PCC 7367, do not encode a FeoB protein for Fe(II) uptake. This study has expanded on the Kranzler *et al*., (2014) data set by searching for genes encoding the non-specific metal ion transporters; the zinc-iron permease (ZIP) and the natural resistance associated macrophage protein homologue (NRAMP), the Fe(II) transporter, FeoB, and the Fe(III) transporters; FutABC and the iron permease, cFTR1, in the genomes of 125 Cyanobacteria, including *Synechococcus* sp. PCC7336, as well as siderophore associated uptake genes. The expression of *cftr1*, the cytochrome c oxidase gene, *cyoC*, and the intracellular iron transcriptional regulator gene, *furA*, in cultures of *Pseudanabaena* sp. PCC7367 grown in an anoxic atmosphere with 0.2% CO_2_ was also investigated. Additionally, we employ a sequence-based approach (using phylogenetic and Bayesian molecular clock analyses) to estimate when iron specific transporters for Fe(II) (namely FeoB) and Fe(III) (namely FutB and cFTR1) appeared within the evolutionary history of the Cyanobacterial Phylum.

## Results

### Iron transporters of *Pseudanabaena* sp. PCC7367 and other basal Cyanobacteria

Initial similarity searches for a FeoB homologue in *Pseudanabaena* sp. PCC7367 indicated that this strain encodes neither the Fe(II) transporter, FeoB, (Supp. Table 1; Kranzler *et al*., 2014 - SI Fig. 4), nor homologues for the standard ZIP and NRAMP metal ion transporters. Instead, it carries genes for Fe(III) specific uptake via FutB, and the reductive iron uptake transporter, cFTR1 (Fig. 2B). Similarly, the basal marine cyanobacterium, *Synechococcus* sp. PCC 7336, (Sánchez-Baracaldo 2015; Boden *et al*., 2021), which is incapable of surviving under a simulated anoxic ferruginous ocean (Herrmann *et al*., 2021), also encoded neither FeoB, nor an NRAMP homologue (Fig. 2B). Expanding the similarity searches for iron transporters identified that most basal Cyanobacteria do not encode FeoB transporters, as it is only encoded in five of sixteen genomes, including *Acaryochloris* spp., *Cyanothece* sp. PCC7425, *Thermosynechococcus elongatus* BP1 and *Synechococcus* sp. PCC6312 (Fig. 2B; Supplementary Table 2).

**Fig 2:**
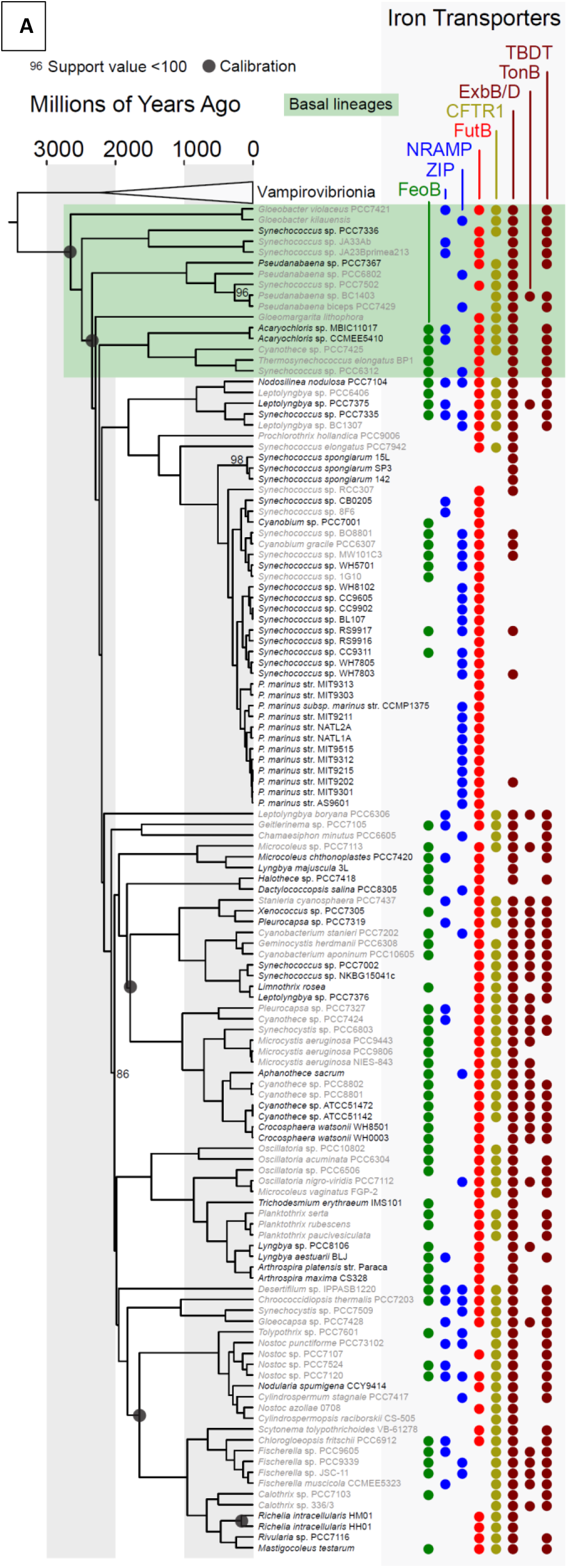
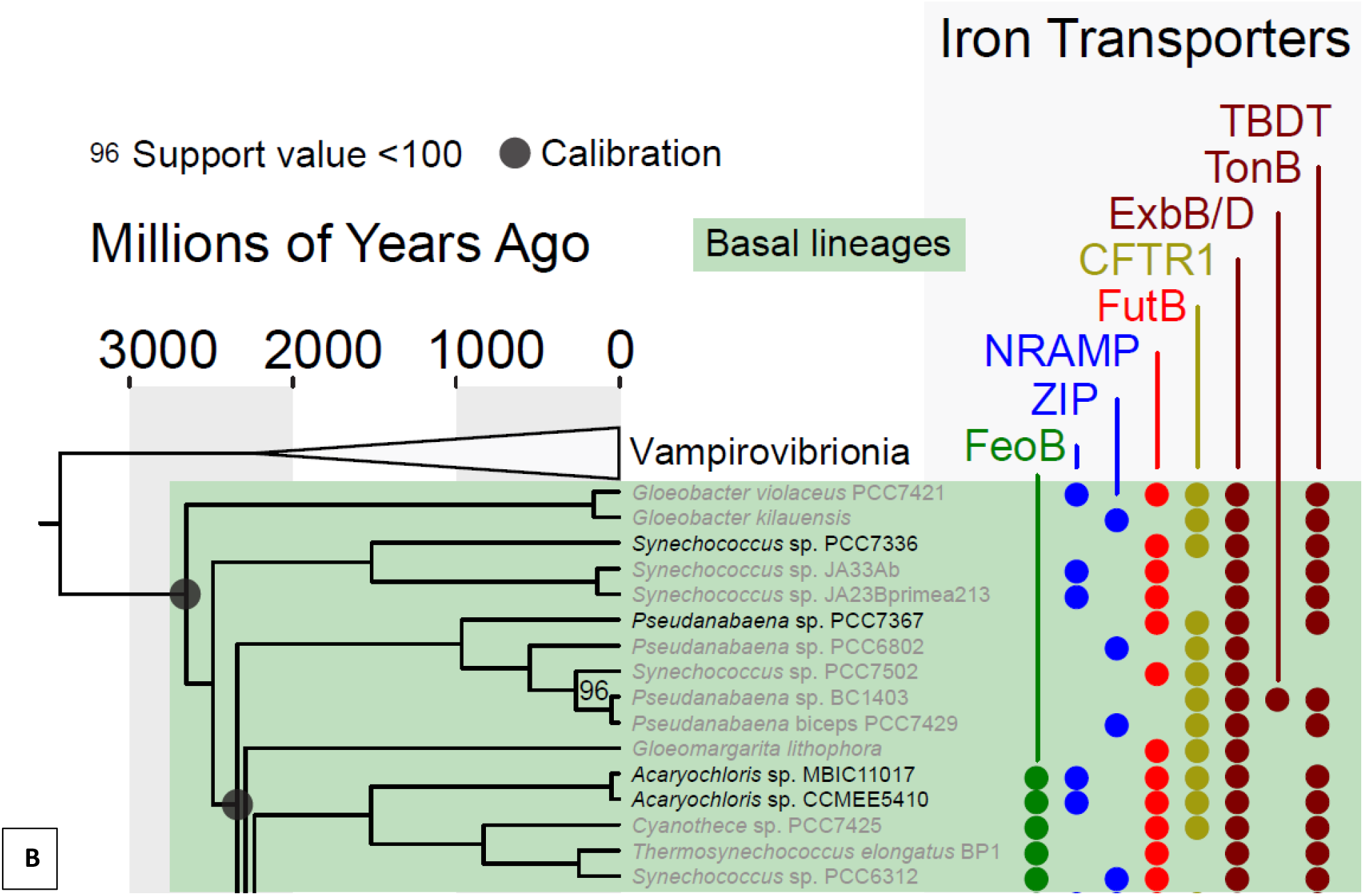
A) Genomic tree of Cyanobacteria indicating the distribution of iron transporters investigated in this study. B) Enlarged section of the genomic Cyanobacterial tree highlighting basal lineages. The inorganic Fe (II) transporters FeoB, ZIP and NRAMP, as well as the Fe (III) transporters, FutB and Cyanobacterial FTR1, are superimposed on a Bayesian molecular clock calibrated tree adapted from (Boden *et al*. 2021). Since the TonB, ExbB/D & TBDT system can also play a role in inorganic iron uptake (Qiu *et al*. 2018) the presence of TonB and TBDTs is also indicated. Support values for branching relationships represent ultrafast bootstrap approximations (Hoang *et al*. 2018). These are equal to 100 unless otherwise stated. The annotations for inorganic iron transporters are as follows: FeoB: green circle; ZIP & NRAMP: blue circles; FutB: red circle; FTR1: olive circle; ExbBD, TonB and TBDTs: brown circles. The names of cyanobacteria isolated from marine habitats are coloured black, in comparison to strains from freshwater, terrestrial and geothermal springs which are grey. Basal lineages (Boden *et al*. 2021; Sánchez-Baracaldo 2015) are indicated inside the green box and enlarged in panel B. Black circles represent calibration points described in Boden *et al*. (2021, Table 1). The first diversification of Cyanobacteria was constrained to occur between 2.32 and 2.7 billion years ago based on evidence of the GOE (Bekker *et al*. 2004) and stromatolitic laminae characteristic of Cyanobacteria (Bosak *et al*. 2009).

In light of the lack of Fe(II) uptake transporters encoded within basal Cyanobacteria, and the potential for the siderophore associated ExbB/D to take up inorganic iron directly (Qiu *et al*., 2018), further similarity searches for siderophore associated uptake genes were conducted. It was confirmed that *Pseudanabaena* sp. PCC 7367 does not produce siderophores (Fresenborg *et al*., 2020; this study), but encodes part of the siderophore uptake system. This includes ExbB/D, and the TonB dependent transporters (TBDTs), *fhuE, iutA* and *fhuA* (Supp. Table 1), but not the TonB protein itself (Fresenborg *et al*., 2020; Årstøl and Hohmann-Marriott 2019). Without TonB, the energy required to transport the siderophore bound Fe(III) through the TBDTs on the outer cell membrane cannot be transferred from the inner membrane proteins, ExbB and ExbD (Årstøl and Hohmann-Marriott 2019; Stevanovic *et al*., 2012). As they are known to be expressed under iron depleted conditions (Fresenborg *et al*., 2020; Årstøl and Hohmann-Marriott 2019), we did not investigate their expression under ferruginous conditions. A similar situation can be observed in other basal Cyanobacteria. All the basal genomes tested contained homologs of ExbB/D and most encoded homologues of TBDTs, but only one basal Cyanobacterium, *Pseudanabaena* sp. BC1403, encodes the accompanying TonB protein (Fig. 2B). *Pseudanabaena* sp. PCC 7367 encodes a few porins that permit the non-specific entry of substances such as metals into the periplasmic space, however their functionalities are not well characterized (Supplementary Table 4). The iron selective porin identified in *Synechocystis* sp. PCC6803 is not present in *Pseudanabaena* sp. PCC7367 (Qiu *et al*., 2021), however 8 potential outer membrane porins were identified (Supplementary Table 4).

### Phylogeny of Cyanobacterial iron uptake genes FeoB, cFTR1 & FutB

In light of the above observations, a broader range of genomes spanning the cyanobacterial tree of life (specifically all of those analyzed in Boden *et al*., 2021) were screened for the presence of iron transporters (Fig 2A, Supplementary Table 2). Genes encoding FeoB, FutB and cFTR1 were found in a variety of strains from marine and non-marine habitats (Fig, 2A), so Bayesian protein phylogenies were generated to describe how these iron transport proteins from different strains of Cyanobacteria are related.

*Thermosynechococcus elongatus* BP-1 is one of five basal Cyanobacteria genetically capable of synthesizing FeoB (Fig 2B). Its genome encodes two FeoB proteins which are distantly related to each other. One of them (NP_682238.1) shares its most recent evolutionary history with the FeoB proteins of *Oscillatoria* sp. PCC6506, *Desertifilum* sp. IPPASB1220 and *Planktothrix serta* (PP 74), whereas the other shares its most recent evolutionary history with different strains, including nitrogen-fixers (e.g. *Fischerella* spp., *Cyanothece* sp. ATCC 51472 and *Trichodesmium erythreaum* IMS101) and unicellular species, such as *Synechocystis* sp. PCC6803 (PP 100) (Supplementary Figure 1). This lack of relationship between the FeoB homologs of *Thermosynechococcus* suggests that its two FeoB sequences have different evolutionary origins.

In contrast to FeoB, which was encoded in the genomes of only five of sixteen basal strains, the ferric iron transporter, FutB, was identified in twelve, the majority of basal strains (Fig. 2). Some of these FutB sequences from basal lineages are closely related. For example, the FutB proteins of *Thermosynechococcus elongatus* BP1 and *Synechococcus* sp. PCC6312 are sisters (PP 100). Similarly, FutB proteins of *Gloeomargarita lithophora, Synechococcus* sp. JA33Ab and *Synechococcus* sp. JA23B’a213 are monophyletic (being derived from a single common ancestor which did not give rise to any other FutB proteins; PP 100, Supplementary Figure 2, Fig. 5).

Unusually, the FutB transporter sequence in *Gloeobacter violaceus* sp. PCC 7421 is removed in an indeterminate group far from those of other basal lineages (PP 100). The FutB transporter sequences for the picocyanobacteria also form a distinctive grouping (PP 100), indicative of a separate origin or high degree of divergence from other Cyanobacterial FutB proteins.

Similar to FutB, cFTR1 proteins are encoded in the genomes of most basal Cyanobacteria (Fig. 2). This includes two *Gloeobacter* spp., which separated from other Cyanobacteria more than 2 billion years ago (Boden et a, 2021). Their FutB homologs are related to those of some other basal lineages, such as *Pseudanabaena* spp., as well as more diverged lineages including *Leptolyngbya boryana* PCC6306 and *Chamaesiphon minutus* PCC6605 (PP 78, Supplementary Figure 3). Like the FeoB paralogs described above, there are also paralogues of cFTR1 in some basal species, such as *Pseudanabaena* sp. PCC 6802 and *Pseudanabaena* sp. BC1403 (Supplementary figure 3). When two homologs of a single protein are found in a single genome, it is possible for one of those homologs to evolve a new and novel function.

### Dating the Cyanobacterial iron transporters FeoB, FutB and cFTR1

The differing distributions of FeoB, FutB and cFTR1 proteins among Cyanobacteria could reflect differences in each strain’s metal requirements and environmental history. We therefore searched for congruence between the evolutionary history of each protein and an established molecular clock (Boden *et al*., 2021) to determine approximately when the *feoB, futB* and *cftr1* genes were introduced into ancestral Cyanobacteria. Similar methods have been utilized previously to map the origin of oxygen-utilizing enzymes in bacteria (Jabłońska and Tawfik, 2021) and predict which genes were present in the last universal common ancestor of life (Weiss *et al*., 2016). It is based on the premise that genes inherited vertically from parental lineages have the same evolutionary history as the species they are found within. A more detailed discussion of how we account for horizontal gene transfer is presented in the Supplementary Information.

We found no phylogenetic evidence that genes encoding the Fe(II) transporter, FeoB nor the Fe(III) transporters, FutB and cFTR1, had been inherited from the ancient common ancestor (specifically the most recent common ancestor (MRCA)) of all extant cyanobacteria. Instead, congruence between the Bayesian molecular clock of Boden *et al*, (2021) and the evolutionary history of FeoB indicate that this ferrous iron transporter traces back to 661 Mya (confidence intervals (CI) range from 1680 to 237 Mya; Table 1) in the Cyanobacterial lineage, which gives rise to *Trichodesmium erythreaum* IMS1 and *Lyngbya aestuarii* BLJ (Figure 3). An earlier origin of FeoB would be possible if the sister homologs found in *Limnothrix rosea* and *Halothece* sp. PCC 7418 (PP 100) were inherited from their common ancestor, which diverged ∼1830 Mya (CIs range from 1954 to 1723 Mya; Table 1).

**Table 1:** Posterior age estimates timing the diversification of ancestral Cyanobacteria with genes encoding FeoB, FutB and CFTR1. ‘Upper’ and ‘Lower’ refer to confidence intervals estimated by the Bayesian molecular clock of Boden *et al*. (2021). MRCA: Most recent common ancestor.

| Fe Transporter | Estimated Age / Mya |  |  | Strains whose MRCA contained the transporter (based on matching evolutionary histories between the transporter and strains) |
| --- | --- | --- | --- | --- |
|  | Mean | Upper | Lower |  |
| FeoB | 661 | 1680 | 237 | *Trichodesmium erythraeum* IMS101<br>*Lyngbya aestuarii* BLJ<br>*Lyngbya* sp. PCC 8106 |
| FutB | 1022 | 1758 | 388 | *Microcystis aeruginosa* NIES 843<br>*Microcystis aeruginosa* PCC 9443<br>*Microcystis aeruginosa* PCC 9806<br>*Cyanothece* sp. PCC 7424 |
| cFTR1 | 1822 | 1965 | 1700 | *Fischerella muscicola* CCMEE 5323<br>*Fischerella* sp. PCC 9339<br>*Chlorogloeopsis fritschii* PCC 6912<br>*Fischerella* sp. PCC 9605<br>*Rivularia* sp. PCC 7116<br>*Mastigocoleus testarum*<br>*Scytonema tolypothrichoides* VB 61278<br>*Calothrix* sp. 3363<br>*Calothrix* sp. 3363<br>*Calothrix* PCC 7103<br>*Chroococcidiopsis thermalis* PCC 7203<br>*Gloeocapsa* sp. PCC 7428<br>*Synechocystis* sp. PCC 7509 |

**Figure 3.**
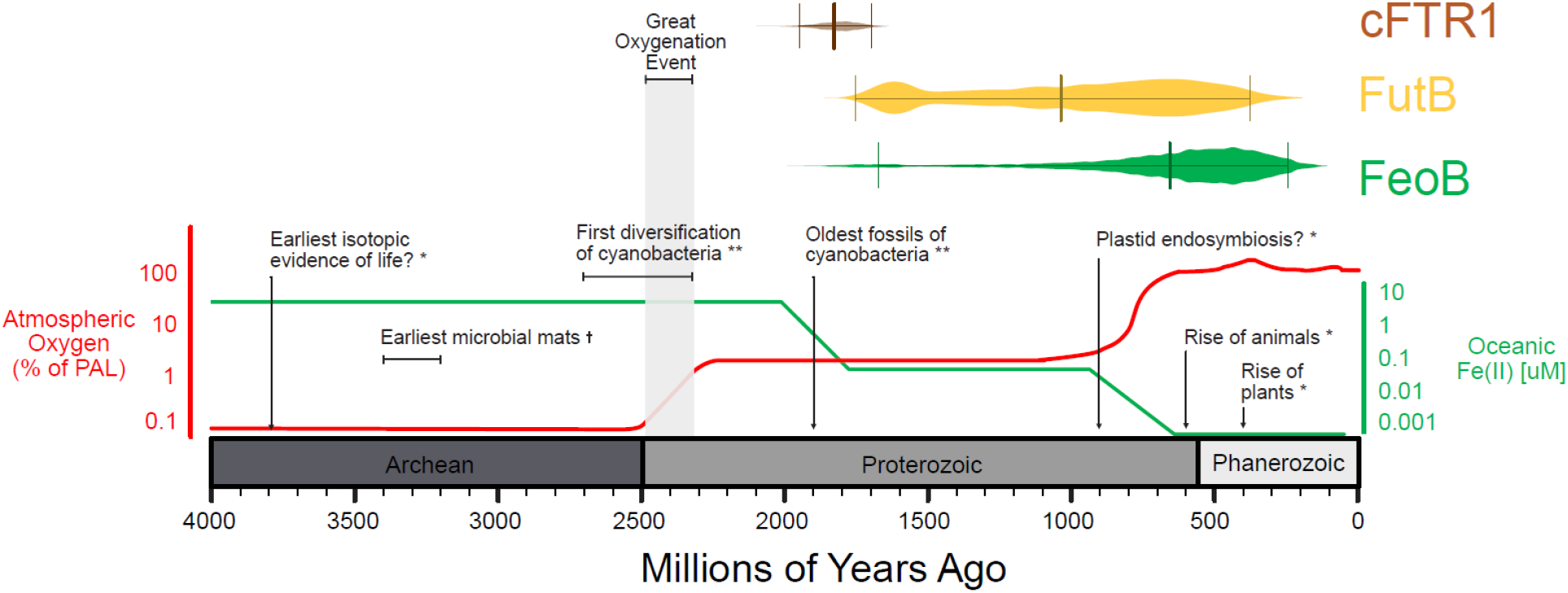
Emergence of Cyanobacterial iron transporters in relation to the oxygen record. The dates for the emergence of the proteins FeoB (Fe(II)) and FutB (Fe(III)) are superimposed on a plot of atmospheric oxygen content (Figure modified from Shih 2015) plotted as the percentage present atmospheric levels (PAL) of O_2_. The dating is based on a Bayesian molecular clock of Cyanobacteria (Boden *et al*. 2021). The Archean ocean Fe(II) levels as calculated by (Saito *et al*. 2003) are plotted against the right hand Y axis. * Shih 2015, †Djokic *et al*. 2021, ** Sánchez-Baracaldo and Cardona 2020; Boden *et al*., 2021.

The high efficiency ferric uptake transporter FutB may have appeared in Cyanobacteria late in the Proterozoic era, at ∼ 1022 Mya (CIs span 1758 to 387 Mya) (Figure 3). This is based on the relationship of FutB homologs found in the freshwater strains *Microcystis aeruginosa* and *Cyanothece* sp. PCC 7424. Their FutB homologs are sisters (PP 100, Supplementary Figure 2), as would be expected if they had been inherited from the strain’s MRCA. It is notable that the basal strain, *Pseudanabaena* sp. PCC 7367, seems to have provided the FutB protein that diversified into unicellular diazotrophs and related Cyanobacteria (PP 80).

The other Fe(III) associated iron transporter, cFTR1 was also present in the Proterozoic when the MRCA of *Chroococcidiopsis thermalis* PC7203 and heterocyst-formers, such as *Nostoc* sp. PCC 7120, radiated ∼ 1822 Mya (CIs range from 1965 to 1700 Mya; Table 1). An earlier origin of cFTR1 could be possible because the homologs found in *Gloeobacter* spp. are sister to one present in *Pseudanabaena* sp. BC1403. If cFTR1 was present in the crown Cyanobacteria and inherited by *Pseudanabaena* sp. BC1403 and *Gloeobacter* spp., but lost in all other lineages, then this topology could be reminiscent of an ancestral cFTR1 present in Cyanobacteria in the Archean. However, a single horizontal gene transfer event between the *Gloeobacter* lineage and *Pseudanabaena* sp. BC1401 could create the same pattern with a more recent origin, so we can only conclude with certainty that cFTR1 was present in the Proterozoic.

### Expression *cFTR1* & *cyoC* in a simulated Archean atmosphere

Previous studies suggested that *Pseudanabaena* sp. PCC7367 was able to withstand nightly influxes of Fe(II) in the Archean ocean, whereas the *Synechococcus* sp. PCC7336 could not (Herrmann *et al*., 2021). Our bioinformatic investigations indicated that the FeoB transporter, responsible for Fe(II) uptake, was encoded in neither species’ genome. *Pseudanabaena* sp. PCC7367 did, however, encode the FutB and cFTR1 transporters involved in Fe(III) uptake. Expression of *futB* is known to be constitutive and not influenced by Fe(II) availability (Katoh *et al*., 2001). In contrast, the cFTR1 transporter was demonstrated to preferentially take up Fe(III) in *Synechocystis*, with an increase in its expression observed under iron starvation (Xu *et al*., 2016). As no alternative respiratory terminal oxidase (ARTO) was identified in *Pseudanabaena* sp. PCC7367 (Fig. 1), we decided to investigate the expression of the gene encoding another terminal oxidase, cytochrome *c* oxidase to ascertain whether it is possibly influenced by the redox state of environmental iron (Schmetterer, 2016). While normally responsible for generating a proton gradient across the thylakoid membrane through the formation of water in the cytoplasm during respiration, cytochrome *c* oxidase may also be involved in modulating the Fe(II)/Fe(III) pool in the periplasm during respiration (Schmetterer 2016). If it is involved in Fe(III) reduction, we would expect a decrease in its expression after the addition of Fe(II). Changes in expression of *cftr1* and *cyoC* were monitored in response to an evening influx of Fe(II) under anoxic conditions.

The Fe(II) and oxygen levels in the media were assessed over 14 hours and are presented in Figure 4. When oxygen levels dropped to zero, one hour after dark, with gentle agitation of the cultures, Fe(II) was added (Figure 4) and its level tracked using the ferrozine assay (Figure 4). Fe(II) was gradually oxidised or taken up overnight, but was still present at 50 µM when the lights went on, after which it was rapidly oxidised as oxygenic photosynthesis commenced.

**Fig. 4.**
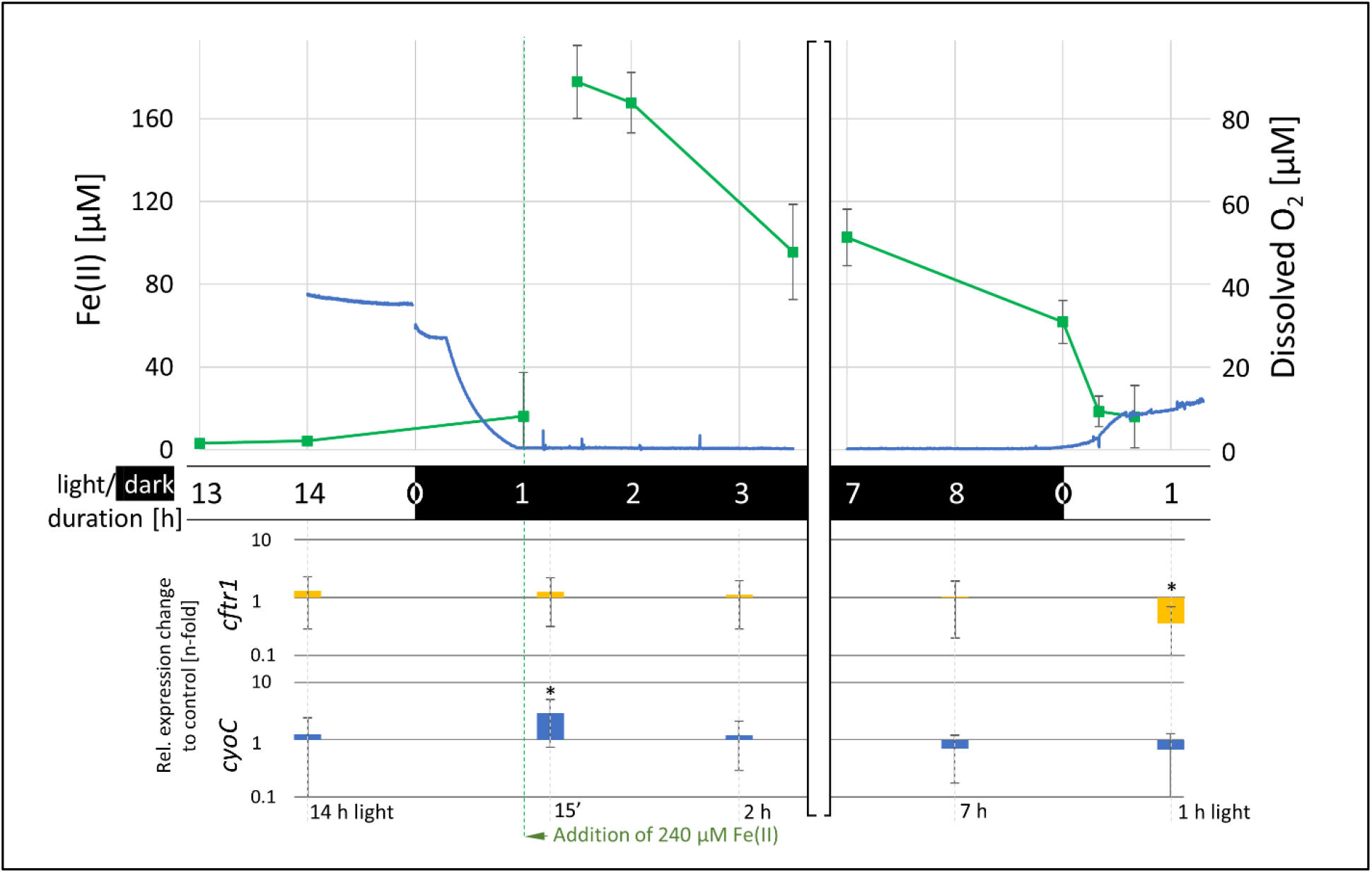
Concentrations of dissolved oxygen and Fe(II) with relative *cftr1* and *cyoC* gene expression levels over the course of a night cycle. The concentration of dissolved oxygen in the culture (blue line), was tracked using an oxygen microsensor, while the Fe(II) concentrations (green line) were periodically determined using the ferrozine assay. The first sample for RNA extraction was taken 1 hour before the dark cycle when oxygen was still present in the medium. The following samples for RNA extraction were taken 15 min, 45 min, 2 h and 7 h after the addition of Fe(II) to a concentration of 240 µM in the dark (green dotted line), when dissolved oxygen levels reached zero. The last sample for RNA extraction was taken one hour after the start of the light cycle. The relative expression of *cftr1*and *cyoC* in the cultures to which Fe(II) was added, in comparison to expression of the control cultures without Fe(II) addition (Supplementary Figure 6), is plotted at the corresponding time points (n = 3). Stars (*) represent a significant difference to the control culture (p<0.05; Student’s t-test, two-tailed).

The relative expression data shows a stable expression of *cftr1* slightly higher than for the control throughout the experiment, with a significant decrease relative to the control an hour after the lights went one. This decrease corresponds to a rapid decrease in Fe(II) in the medium (Figure 4). The expression of cytochrome c oxygenase increased significantly after the addition of Fe(II), decreasing to late daytime levels two hours after the addition of Fe(II).

## Discussion

The GOE was one of the most significant developments in Earth’s history, because in a relatively short period of geological time, the atmosphere changed from an anoxic reducing environment to an oxic one. Significantly, soluble Fe(II) was oxidized to barely soluble Fe(III) in the photosynthetically active top layers of aquatic environments, suggesting that early life had to evolve means to overcome iron limitation. Our hypothesis was that, given a continuous lineage, the handful of basal taxa of ancient Cyanobacteria that left modern-day descendants would have retained some ancestral features (Raven and Sánchez-Baracaldo, 2021), with evidence of FeoB, FTR1, TonB and ExbB/D in their genetic complement prior to the GOE (Qui *et al*., 2021; Fresenborg *et al*. 2020). FeoB, is considered the primary Fe(II) transporter in Cyanobacteria, based on studies of the Cyanobacterial model organism, *Synechocystis* sp. PCC 6803, which preferentially takes up Fe(II), rather than Fe(III) via FutB (Katoh *et al*., 2001). Previous investigations highlighted differences in growth responses between two of marine basal strains of Cyanobacteria, *Pseudanabaena* sp. PCC 7367 and *Synechococcus* sp. PCC 7336, in response to a tidal influx of Fe(II) seawater at night (Herrmann *et al*., 2021). We therefore wanted to compare the genetic potential of these strains with respect to iron uptake transporters, as well as investigate the expression of the cFTR1 Fe(III) uptake associated gene in *Pseudanabaena* sp. PCC7367 after a tidal influx of Fe(II) in a simulated Archean shallow water marine environment.

Similarity searches to identify FeoB in 125 Cyanobacterial genomes, identified a general absence of genes encoding FeoB in basal Cyanobacteria (Fig. 2), with it only identified in five strains. As a result, FeoB may be the dominant iron transporter in in the more recently diverged *Synechocystis* sp. PCC 6803 (Katoh *et al*., 2001), but appears not to be representative of all Cyanobacteria, especially the basal lineages. Specifically, both *Pseudanabaena* sp. PCC 7367 and *Synechococcus* sp. PCC7336, as well as the previously investigated *Synechococcus* sp. PCC7002, do not encode FeoB transporters. Neither do these strains encode the iron permease, ZIP, nor NRAMP (Fig. 2), thereby raising the question as to how they would have taken up Fe(II) from the ferruginous Archean ocean. The presence of 8 cyanobacterial putative outer membrane porins encoded on the genome of *Pseudanabaena* sp. PCC7367 (Supplementary Table 4) potentially provide a means for inorganic iron and other metals to enter the periplasmic space for transport across the cell membrane.

Localized oxidation of Fe(II) during the day would have provided Fe(III) in the vicinity of *Pseudanabaena* sp. PCC 7367 (Herrmann *et al*., 2021), suggesting that Fe(III) may have been the preferred source of iron. Both *Pseudanabaena* sp. PCC 7367 and *Synechococcus* sp. PCC7336 encode the Fe(III) transporters, FutABC, cFTR1, as well as the ExbB/D and TBDTs, without the energizing TonB component (Fig. 2) and do not encode siderophores (Årstøl and Hohmann-Marriott 2019; Fresenborg *et al*. 2020). This is in contrast to the previously studied derived *Synechococcus* sp. PCC7002 (Swanner *et al*., 2015a; Swanner *et al*., 2015b), which emerged in the phanerozoic (Sanchez-Baracaldo, 2015) and, in addition to FutB, cFTR1, ExbB/D and TBDTs, encodes a functional TonB as well as synthesizing siderophores (Fig. 2, Supplementary Table 2) (Årstøl and Hohmann-Marriott 2019; Fresenborg *et al*., 2020), suggesting Fe(III) is its primary source of iron.

Given how essential Fe is for cyanobacteria and how its availability has changed over geological time, we decided to investigate the evolutionary history of FeoB, FutB and cFTR1 within Cyanobacteria. Bayesian trees indicated a complex history involving gene duplication and/or other patterns of reticulated evolution, leading to individual strains sometimes harbouring more than one gene for a given transporter. For example, two FeoB homologues with different evolutionary trajectories were found in the deeply branching *Thermosynechococcus elongatus* BP1 (Supplementary Figure 1). Furthermore, the FutB homologue of the cyanobacterium, *Gloeobacter violaceus* sp. PCC 7421 was unrelated to FutB homologs of other basal strains (Supplementary Figure 2).

Given this scattered phylogeny of iron transporters in basal lineages, the evolutionary history of each iron transporter was compared to an existing genome tree of Cyanobacteria (presented in Boden *et al*., 2021) to search for congruence indicative of inheritance from a common ancestor. Further comparisons to a previously published molecular clock (Boden *et al*., 2021) facilitated an estimation timing the appearance of each transporter. Overall, this revealed that the Fe(II) transporter, FeoB, and the Fe(III) transporters, FutB and cFTR1 were present in the Cyanobacterial lineage during the Proterozoic (Fig. 4) when atmospheric O_2_ levels were ∼1% of present day levels. If one were to trace the predictions of oceanic iron levels in the ocean (Saito *et al*., 2003), one would see that lineages with FeoB appeared more widely during the Neoproteozoic Oxygenation Event, when the Fe(II) levels dropped to ∼1 nM in the open ocean, whereas lineages with the Fe(III) transporters, FutB and cFTR1, were likely already present (Fig. 4). Our phylogenetic analyses, although limited by the availability of Cyanobacterial genome sequences (future discovery of novel lineages, particularly of basal strains, could change our predictions) also fail to find evidence that FeoB, FutB and cFTR1 were present in Cyanobacteria during the Archean, suggesting they were incorporated into Cyanobacterial genomes later, when Fe(II) levels dropped in the oceans.

Expression of the iron permease gene, *cftr1*, in the control cultures of *Pseudanabaena* sp. PCC 7367 remained largely unchanged during the complete 24-hour cycle, with a non-significant increase in expression recorded an hour after the lights went on and oxygen levels rose. This upwards trend augmented the reduced relative expression of *cftr1* in the experimental cultures exposed to Fe(II) after the start of the light cycle (Fig. 4), suggesting a role for O_2_ and iron speciation in the regulation of *cftr1* expression.

This is the first time that *cftr1* transcription has been confirmed in a non-*Synechocystis* species, and under anoxic, ferruginous conditions. However, the expression data contradicts that observed in *Synechocystis*, whereby *cftr1* expression was induced by iron starvation (Xu *et al*., 2016). Additionally, the diurnal cycling of the expression of metal transporters as recorded in *Synechocystis* sp. PCC6803 (Saha *et al*., 2016), was not observed in our study. Expression of diurnally regulated genes, including some involved in metal transport, were upregulated in *Synechocystis* sp. PCC6803 an hour before and two hours into the light cycle (Saha *et al*., 2016). As we sampled two hours prior to the light cycle, we may not have recorded diurnally induced increased expression at the end of the dark cycle. Further investigations regulating iron speciation and uptake is required to better understand the process of iron acquisition in *Pseudanabaena* sp. PCC7367 and other basal Cyanobacteria under iron replete conditions.

ARTOs have been proposed to provide reduced Fe(III) to the cFTR1 permease in *Synechocystis* sp. PCC 6803 (Kranzler *et al*., 2014). While *Synechococcus* sp. PCC 7336 encodes an ARTO homologue, *Pseudanabaena* sp. PCC 7367 does not (Schmetterer 2016, this study). As there is some evidence for cytochrome c oxidase to be located on the cytoplasmic membrane in *Trichodesmium thiebautii* (Bergman *et al*., 1993), we decided to investigate its expression in *Pseudanabaena* sp. PCC 7367 after the addition of Fe(II) (Fig. 4). Terminal oxidase expression is known to be increased at night during respiration, regulated by the circadian clock in *Synechococcus elongatus* PCC7942 (Ito *et al*., 2009) and the iron uptake regulator, FurA, in *Anabaena* sp. PCC7120 (González *et al*., 2012; González *et al*., 2014). Interestingly, expression of the cytochrome c oxidase increased significantly after the addition of Fe(II) before dropping to its original level two hours after Fe(II) addition.

As pre-GOE Cyanobacterial lineages have a continual lineage with present-day strains, it would imply early species could not access the Fe(II) in their localized environments. In fact, with the late incorporation of FutB, FeoB and cFTR1 into present-day Cyanobacterial lineages it would appear that the cation transporters ZIP and NRAMP may have been the only active Fe(II) transporters available for most basal Cyanobacteria during the Archean, however neither is encoded by *Pseudanabaena* sp. PCC7367 (Fig. 2) (Qui *et al*., 2021; Fresenborg *et al*., 2020). This would suggest that the Archaen Cyanobacteria may have been iron limited, thereby reducing their growth rates.

Siderophore synthesis is also not a feature of the basal clade of Cyanobacteria, although most strains carry genes encoding siderophore uptake transporters (TBDTs) into the periplasmic space (Årstøl and Hohmann-Marriott 2019; this study Fig. 2). Fe(II) can be mobilized from mineral sources by siderophores, specifically desferroxazine (Kraemer, 2004; Bau *et al*., 2013), produced by several micro-organisms in niches with low iron availability. This siderophore is also able to be transported by the non-siderophore producing *Synechocystis* sp. PC6803, suggesting that scavenging of siderophore bound Fe(III) may have been the initial mode by which Cyanobacteria were able to meet their iron requirements during the Archean. While TBDTs and the ExbB/D proteins are largely present in the basal clade Cyanoabcterial strains, the TonB protein is not encoded, suggesting that some other mechanism may exist to power siderophore:Fe(III) complexes through the outer membrane into the periplasmic space (Årstøl and Hohmann-Marriott 2019). It has been suggested that the ExbB/D siderophore transporter, also encoded by *Pseudanabaena* sp. PCC7367, is capable of mineral Fe(III) uptake in *Synechocystis* sp. PCC6803 (Qiu *et al*. 2018). The high levels of oxygen measured in freshwater mats of *Gloeobacter violaceus* PCC7421 and *Chroococcidiopsis thermalis* PCC7203 (Herrmann & Gehringer, 2019), as well as that recorded in liquid cultures of *Pseudanabaena* sp. PCC7367 (Herrmann *et al*., 2021) in anoxic atmospheres suggest that there would have been sufficient Fe(III) available for putative ExbB/D uptake in the immediate vicinity within these oxygen rich niches. Future studies can investigate siderophore mobilization of Fe(II)/Fe(III) under anoxic conditions, both in the presence and absence of siderophores, as well as conducting additional molecular analyses to date the inclusion of ExbB/D in the Cyanobacterial phylum. The presence of other, as yet unidentified iron chelators in Cyanobacteria cannot be excluded, as evidenced by the recent identification of Cyanochelins in some Cyanobacterial species (Galica *et al*., 2021). Cyanochelin gene cluster was identified in a single basal species, namely *Synechococcus* sp. PCC7336, and only induced under iron depleted conditions (Galica *et al*., 2021).

The mechanisms by which ancestral Cyanobacteria acquired Fe(II) / Fe(III) in the ferruginous Archean environment, remains unclear. Under the iron limiting conditions of today’s ocean, FutB must play an essential role in iron acquisition in *Pseudanabaena* sp. PCC 7367, however confirmation of its constitutive expression, as reported for *Synechocystis* sp. PCC 6803 (Katoh *et al*. 2001), especially under conditions of high Fe(II) availability, is required. In conclusion, this study has challenged the accepted viewpoint that Cyanobacteria had unlimited access to Fe(II) in the Archean ocean, as the majority of basal strain descendants that survived the Archean do not encode the necessary Fe(II) transporters. Furthermore, this study raises questions as to the influence of oceanic trace metal inventories on the evolution of biochemical pathways, particularly with respect to metalloenzymes (Dupont *et al*., 2006 & 2010). Given the recent support for the evolution of oxygenic photosynthesis close to the origin of life (Oliver *et al*., 2021; Fournier *et al*., 2021), basal Cyanobacteria may have been utilizing Fe(III), from their inception, via the TBDT and ExbB/D system to meet their high iron requirements, while possibly scavenging siderophore bound Fe(III) from other community members. The data presented here has highlighted the need to further investigate iron uptake by Cyanobacteria, especially under the anoxic, ferruginous conditions on early Earth, in order to obtain a clearer picture of limitations on Cyanobacterial expansion prior to the GOE.

## Materials and Methods

### Gene screening

In order to understand the differences in the perceived Fe(II) toxicity between *Pseudanabaena* sp. PCC7367 and *Synechococcus* sp. PCC7336 (Herrmann *et al*., 2021), it was necessary to identify all iron transporters in these, and other basal lineages of Cyanobacterial. To do this, 125 genomes (Supplementary data file 1) were screened with protein sequence similarity searches (March 2020 & May 2021) using BLASTP (Altschul 1991; Altschul 1993; Zhang *et al*., 2000) with e-value cut-offs less than 0.001 for the following genes linked to iron transporters identified in *Synechocystis* sp. PCC6803, namely FutA, FutB and FutC (Brandt *et al*., 2009; Katoh *et al*., 2001), FTR1 (Katoh *et al*., 2000), FeoB (Katoh *et al*., 2001; Kranzler *et al*., 2014), ExbB/D TonB (Qiu *et al*., 2018; Jiang *et al*., 2015) and the related *E. coli* iron transporters (Altschul, 1991; Altschul, 1993; Zhang *et al*., 2000). The identification of hits found in *Pseudanabaena* sp. PCC 7367 and other basal cyanobacterial lineages (Sánchez-Baracaldo, 2015) was verified with FeGenie (Garber *et al*., 2020). The complete bioinformatics search and processing pipeline is illustrated graphically in Supplementary Figure 7.

### Phylogenetic Analyses

Our initial screen indicated a lack of *feoB* in most basal Cyanobacterial genomes, so the abovementioned gene screen was extended to search for iron transporters in a broader range of genomes representing the full diversity of Cyanobacteria (Boden *et al*., 2021). Phylogenetic analyses were then employed to investigate how iron transporters evolved in Cyanobacteria. To do this, amino acid sequences of FeoB, FutB and Cyanobacterial FTR1 were aligned using MUSCLE (Edgar, 2004) implemented in MegaX version 10.1.8 (Kumar *et al*., 2018) with the following parameters; gap open -2.9, gap extend 0, hydrophobicity multiplier 1.2, maximum iterations 16, cluster method UPGMA, minimum dialogue length 24. Poorly aligned regions, specifically those with more than 80% gap regions, were removed manually. In order to reconstruct the phylogeny of iron uptake genes, Bayesian phylogenetic trees were generated for FeoB, FutB and cFTR1 in MrBayes 3.2.7a (Ronquist *et al*., 2012) using a mixed amino acid substitution model prior, invariant sites and 4 categories of a gamma distribution to model changes in substitution rates across different sites of the alignment. Convergence was assessed for two replicate chains using a burn-in of 25%. When the average standard deviation of split frequencies (ASDSF) was ≤ 0.03, potential scale reduction factors (PSRF) between 1.00 and 1.02 and ESS scores assessed in Tracer v1.6 (Rambaut *et al*., 2013) ≤ 200, trees were considered converged.

### Genome tree and Molecular Clock Analyses

To estimate when the iron uptake genes specific for Fe(II), Fe(III) and the cFTR1 permease emerged in Cyanobacteria, the evolutionary history of FeoB, FutB and cFTR1 were compared to the maximum likelihood phylogeny of Boden *et al*., 2021. Details of how this phylogeny was produced are present in the original paper (Boden *et al*., 2021), which incorporates information from 139 proteins, 16S rRNA and 23S rRNA collected from >100 strains representing the entire diversity of Cyanobacteria. If topology of this species tree matched the topology of Bayesian phylogenies of FeoB, FutB or cFTRA, generated in the present study, then the MRCA of that clade was assumed to have utilized the protein. To find out when those ancestors diversified, we cross-referenced them to the Bayesian molecular clock of (Boden *et al*. 2021). This was made using information from rRNA (16S and 23S) and 6 soft calibrations from fossils and geological records. For further detail, see (Boden *et al*., 2021).

### Culture conditions and experimental set-up

*Pseudanabaena* sp. PCC7367 (Pasteur Culture Collection, Paris, France), was maintained in the prescribed ASNIII medium and acclimated to the simulated Archean atmosphere in an anoxic chamber atmosphere (GS Glovebox, Germany) of N_2_ gas supplemented with 0,2% CO_2_., 17:9 hr day-night cycle, 65% humidity and 25 Photosynthetic Photon Flux Density (PPFD [µmols photons · m^-2^ · s^-1^]) (Herrmann *et al*., 2021). Triplicate cultures, inoculated at 0.4 µg Chl a · ml^-1^ from late exponential phase cultures were set-up in acid washed, sterilized Fernbach flasks containing 600 ml medium equilibrated at the experimental atmosphere. Chl a determination of cell content is routinely used to monitor Cyanobacterial growth and viability. Briefly, Chl a was extracted from a 1.5 ml culture volume on days 1, 3, 6, 9 and 11. Cell pellets were lysed in 90% (v/v) CaCO_3_ neutralized methanol by bead beating, quantified as described in Herrmann *et al*., (2021) and plotted to generate a growth curve for *Pseudanabaena* sp. PCC7367 (Supplementary Figure 1). On day 10, Fe(III) was added to the cultures to ensure they were not iron depleted. The following day, an oxygen microsensor (Ox200, UNISENSE, Denmark) was installed to monitor the O_2_ levels resulting from oxygenic photosynthesis, in the cultures for the duration of the experiment.

### Spectrophometric ferrozine iron assay

Fe(II) and Fe(III) levels were monitored periodically by means of the spectrophotometric ferrozine iron assay to confim the availability of Fe(II) at night (Herrmann *et al*. 2021). Briefly, the cultures in the anaerobic glovebox (GS Glovebox, Germany) were gently resuspended and 2 × 1 ml culture volume was removed under sterile conditions from each biological replicate, added to 2 ml reaction tubes (Sarstedt, Germany) and the particulate matter immediately pelleted by centrifugation at 14 000 x g (Hermle Z 233 M-2) for 1 minute. A volume of 150 µl of the supernatant was diluted 1:1 in anoxic MilliQ H_2_O, in the anoxic workstation, in a pre-prepared 96 microwell plate, to dilute the Fe (II) concentration to the detection range of the assay. For the determination of particulate Fe (III), the pellet was resuspended in 1 ml of 1 N HCl (Roth) outside the glovebox.

Assay standards for Fe(II) ranging from 150 µM to 2.34 µM FeSO_4_•7H_2_O (Sigma-Aldrich) and Fe(III) FeCl_3_ • 6H_2_O (Sigma-Aldrich) were prepared in 1 M HCl. Volumes of 150 µl of each sample or standard were added to a 50 µl volume of buffered ferrozine solution (50% w/v Ammonium acetate, 0.1% w/v Ferrozine (Disodium-4-[3-pyridin-2-yl-6-(4-sulfonatophenyl)-1,2,4-triazin-5-yl]benzosulfonate (Sigma-Aldrich) in dd, H_2_O). In order to determine the amount of Fe (III) in the medium, 150 µl of the samples were added to 50 µl reducing agent (10% (w/v) hydroxyl hydrochloride in 1 M HCl) and incubated for 30 min. in the dark before the addition of 50 µl buffered ferrozine solution. The OD_562_ was measured (Stookey 1970) after 5 minutes incubation, in a microplate reader (Multiscan FC, ThermoFisher Scientific, USA). The Fe(II) and Fe(III) concentrations were determined off the standard curves (R^2^ = 0,9989 and R^2^ = 0,9997 respectively) for the triplicates samples at the timepoints indicated in Fig. 2 and Supplementary Fig. 2 respectively.

### Primer design and validation

Primers for reverse transcription quantitative PCR (RT qPCR), were designed and validated to detect the following genes of *Pseudanabaena* sp. PCC 7367: the Cyanobacterial iron permease, cFTR1 (Pse7367_Rs12485), the ferric uptake regulator, FurA (Pse7367_Rs06445), the cytochrome *c* oxidase, (Pse7367_Rs00935) and the reference target gene, *rpoC1* (Pse7367_Rs07505), encoding the RNA polymerase gamma subunit (Alexova *et al*. 2011). The primer sequences, PCR product length and primer amplification efficiencies are presented in Supplementary table 3. PCR product integrity was confirmed by melt curve analysis of, using cDNA of *Pseudanabaena* sp. PCC7367 as template.

### RNA extraction and synthesis of copy DNA

In order to track the effect of a tidal influx of Fe(II) on the expression of iron transporters in *Pseudanabaena* sp. PCC7367, the cultures were sampled an hour before darkness and, once the levels of O_2_ reached zero, Fe(II) (FeCl_2_) was added to the experimental cultures to a final initial concentration of 240 µM Fe(II). Further samples for RNA extraction were collected 15 min., 45 min., 2 h and 7 h after the addition of Fe(II), in the dark period, with a final sample was collected one hour after the lights went on. Culture volumes of 45 ml of gently resuspended culture material were decanted into a 50 ml Falcon reagent tube containing 5 ml ice cold stop solution (95% Ethanol: 5% Phenol v/v; Roth, Germany) and gently inverted to prevent further transcription. Cells were pelleted at 4 000 x g for 10 min (Eppendorf 5810R, Germany). Throughout the experiment, the cultures were gently agitated by a magnetic stirrer bar set at 150 rpm to facilitate the release of O_2_ from the culture medium prior to the addition of Fe(II) (Herrmann *et al*., 2021). Pelleted cells were drained and stored at -80°C until RNA extraction.

RNA was extracted from the thawed pellets using the NucleoSpin^®^ RNA Plant Kit (Macherey-Nagel, Germany) according to the manufacturer’s instructions, with a modified cell lysis step (Mironov and Los 2015). The cell pellets were transferred to a sterile 2 ml tube (Sarstedt, Germany) containing 100 mg RNAse-free 0.1 mm silica beads (Biospec, Germany). RA1 buffer was added (350 µl of RA1 buffer per 100 mg pellet) to the pellet, as well as 1% (v/v) β-Mercaptoethanol (2-Mercaptoethanol, ROTH, Germany). The samples were frozen in liquid nitrogen, allowed to thaw, then were disrupted for 90 sec at 6.5 m. s^-1^ (Fastprep FP120, Thermo systems, USA) followed by an additional freeze/thaw step.

The cell lysates were centrifuged for 1 min at 14 000 x g (Hermle Z233-M2, Germany) to pellet the cell debris and the RNA was extracted from the supernatant using the two column-system of the NucleoSpin^®^ RNA Plant Kit (Macherey-Nagel, Germany). DNA removal was ensured by the on-column DNA digestion according to the manufacturer’s description. RNA thus obtained was spectrophotometrically quantified (NanoDrop® Lite, Thermo Scientific, USA) and the quality confirmed by agarose gel electrophoresis, with DNA digestion verified by PCR targeting the housekeeping gene, *rpoC1*.

Extracted and purified RNA was reverse transcribed into first-strand copy DNA (cDNA) using the ProtoScript® II Reverse Transcriptase Kit (NEW ENGLAND BioLabs®Inc, Germany), according to the manufacturer’s instructions, using up to 1 µg of RNA template, in RNAse-free microfuge tubes. After cDNA synthesis, the remaining RNA was degraded by the addition of 10 µl 1 M Tris-EDTA and 100 µl of 0.1 M NaOH and incubated at 95 °C for 10 min. The NaOH was neutralized by the addition of 1 M HCl and the cDNA was purified by a PCR-clean-up using the NucleoSpin^®^ Gel and PCR Clean-up kit (Macherey-Nagel, Germany) according to manufacturer’s instructions. The concentration of the newly synthesized cDNA was measured with a NanoDrop^®^ Lite Spectrophotometer (Thermo Scientific, USA), where after the cDNA was stored at -20 °C until use.

### Quantification of gene expression

The levels of expression of the genes encoding FurA, cFTR1 and cytochrome c oxidase and the housekeeping gene for the gamma subunit of the RNA polymerase, *rpoC1*, were determined via quantitative PCR (qPCR) of the cDNA (Huggett *et al*. 2013; Nolan *et al*. 2013). Each 10 µl reaction was prepared with 5 µl 2x iTaq™ Universal SYBR^®^ Green Supermix, 5 pmol of each primer and 10 ng cDNA template. The volume was adjusted to 10 µl with RNase free water. The reactions were performed in triplicate, on three different days, to evaluate transcript abundance relative to the expression of the housekeeping gene (Lü *et al*., 2018) using a BIORAD CFX Connect™ Real-Time System thermocycler. Cycling for the qPCR was as follows: activation of the polymerase (50 °C for 10 min), followed by initial denaturation at 95 °C for 5 min and 40 cycles of 95 °C for 10 sec, 20 sec at the primer-pair specific annealing temperature and 72 °C for 10 sec, with a final elongation step at 72 °C for 5 min. Primer T_m_ and T_a_, as well as the product size, are listed in Table 1. Product length was verified by melt curve analysis and the relative gene expression was calculated from the mean fold difference of ΔC_q_ values of the three biological replicates for each timepoint (Huggett *et al*., 2013; Guescini *et al*., 2008; Rutledge and Stewart, 2008; Narum, 2006), in Excel (Excel 365, Microsoft, USA).

### Statistical analyses

Statistical analyses were done using the two-tailed, heteroscedastic Student’s t-test (Excel 365, Microsoft, USA) to determine the influence of Fe(II) on gene expression levels.

## Acknowledgements

This project was funded by the German Research Foundation, DFG, Grant number: GE2558/3-1 & GE2558/4-1 awarded to MMG, a University of Bristol Graduate Teaching Scholarship awarded to J.S.B. and a Royal Society University Research Fellowship awarded to P.S.B.

## Author contributions

M.M.G., P.S.B. & A.J.H. conceptualized the project and designed the research experiments. T.C.E.B., J.S.B., K.W.E. and A.J.H. conducted the experiments and analyzed the data. All authors contributed to interpreting the data and writing the manuscript.

## Data availability

The sequence data analyzed in this study are available in the open science framework repository, https://osf.io/7x598/?view_only=715cd38c378446ba8c3f6c924f9be9f5. All other data are included in the published article and its supplementary file.

## Additional information

Supplementary Tables.

Supplementary Figures.

