## Supplemental Figures for "Lack of Fe(II) transporters in basal Cyanobacteria complicates iron uptake in ferruginous Archean oceans"

### Supplementary Figures

Tristan C. Enzingmüller-Bleyl[1§], Joanne Boden[2§], Achim J. Herrmann[1], Katharina W. Ebel[1], Patricia Sanchez-Baracaldo[2], Nicole Frankenberg-Dinkel[1], Michelle M. Gehringer[1*]

[1]Department of Microbiology, Technical University of Kaiserslautern, Kaiserslautern, 67663, Germany.

[2] School of Geographical Sciences, Faculty of Science, University of Bristol, Bristol, BS8 1SS, United Kingdom.

[§] These authors contributed equally to the manuscript.

*

Keywords: Iron uptake, Cyanobacteria, *Pseudanabaena* sp. PCC7367, Archean, evolutionary dating

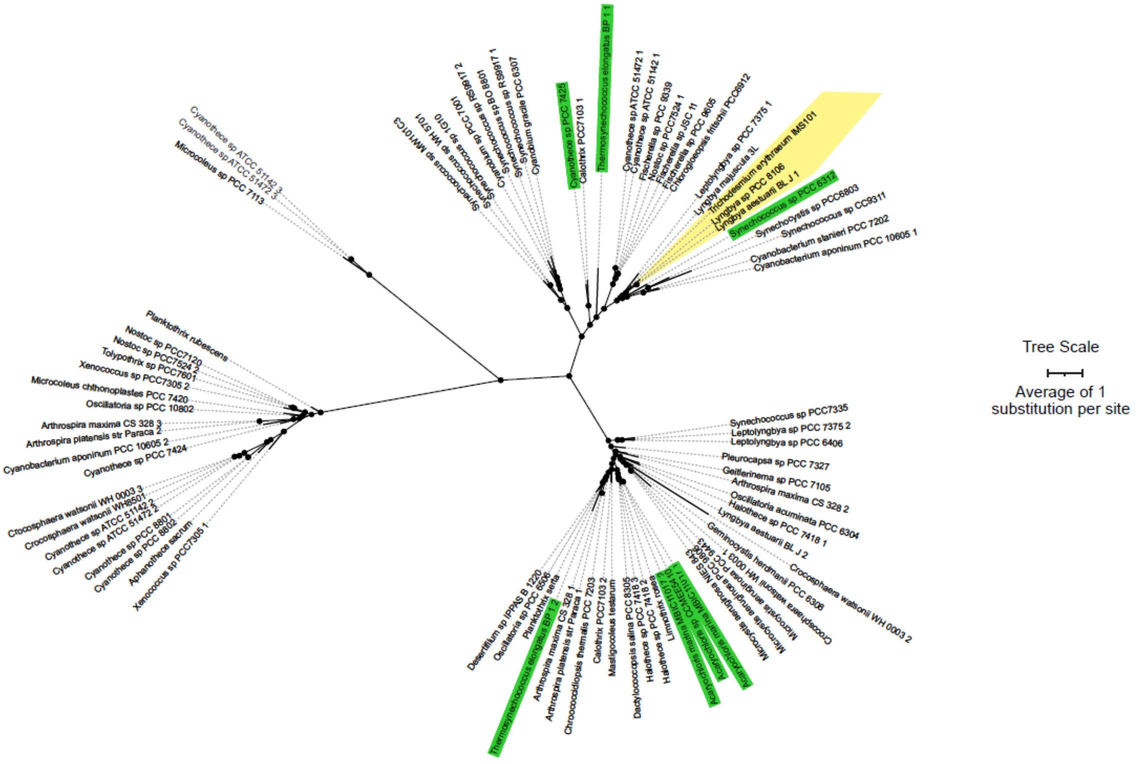


**Supplementary Figure 1. Unrooted Bayesian phylogeny of FeoB.** Protein sequences obtained using BLAST P and the reference gene from *Synechocystis* sp. PCC 6803 (Katoh *et al*. 2001; Kranzler *et al*. 2014) implemented in MegaX (Kumar *et al*. 2018) for 1018 amino acids. The alignment was truncated to 779 amino acids by removing regions of low homology. Phylogenetic reconstruction was performed in MrBayes v3.2.7a (Ronquist *et al*. 2012) using the mixed substitution model option with invariant sites and gamma distributed rates. Taxon labels of basal strains are labelled with a green background for easy identification. Each node in the unrooted tree with a black circle has a posterior probability more than 0.95. Branch lengths represent amino acid substitutions with the scale bar representing an average of 1 substitution per site. Proteins which can be traced back to ancestral cyanobacteria which lived 661 mya are highlighted in yellow.

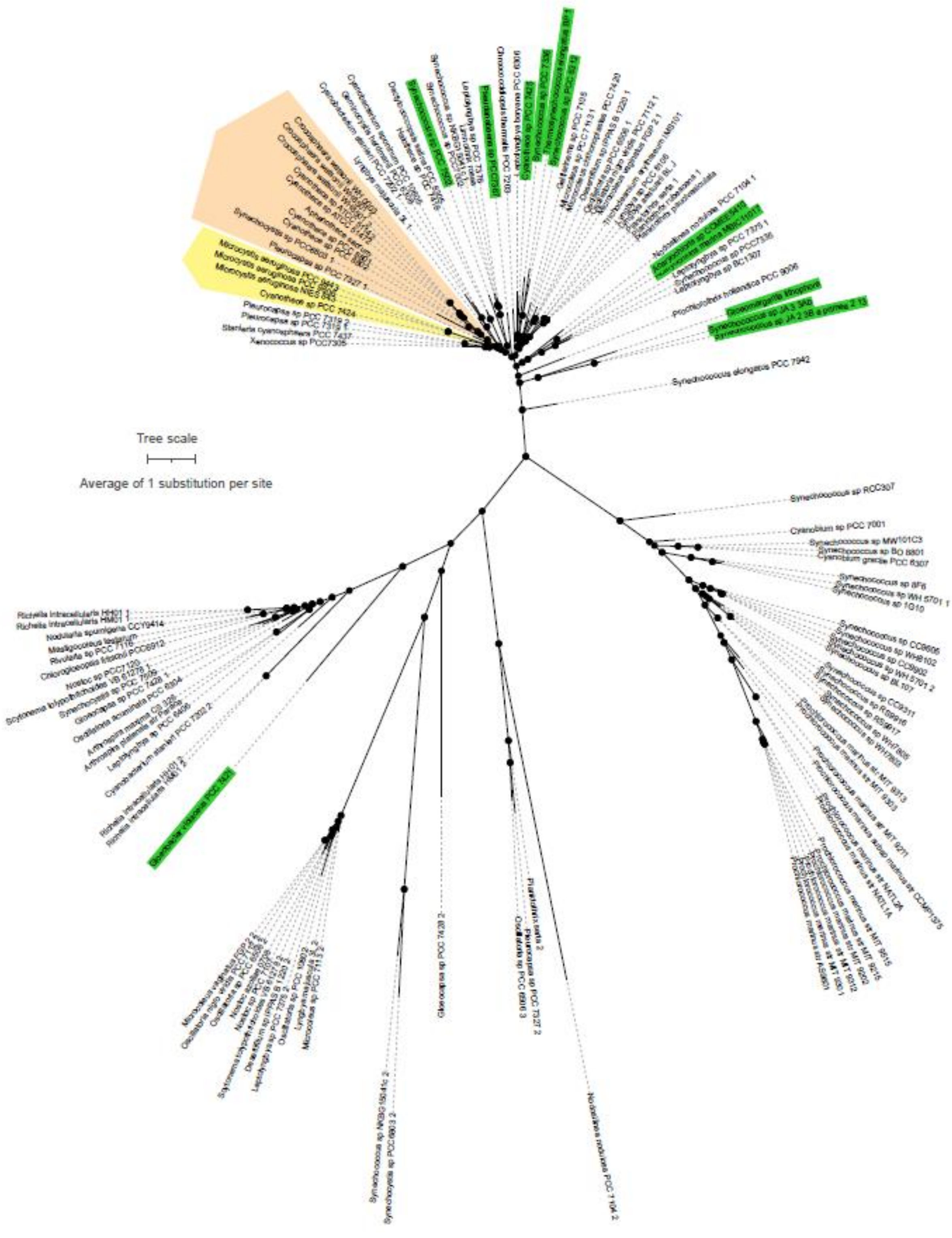
Tree scale
Average of 1 substitution per site
Synechococcus elongatus PCC 7942
Synechococcus sp RCC307
Cyanobium sp PCC 7001
Gloeocapsa sp PCC 7428 2
Nodularia nodulosa PCC 7104 2

**Supplementary Figure 2. Unrooted Bayesian phylogeny of FutB.** Protein sequences obtained using BLASTP and the reference gene from *Synechocystis* sp. PCC 6803 (Brandt *et al.* 2009; Katoh *et al*. 2001) were aligned in MUSCLE (Edgar 2004) implemented in MegaX (Kumar *et al*. 2018). The alignment of ~666 amino acids was truncated to 562 amino acids by removing regions of low homology. Phylogenetic reconstruction was performed in MrBayes v3.2.7a (Ronquist *et al*. 2012) using the mixed substitution model option with invariant sites and gamma distributed rates. Taxon labels of basal strains are labelled with a green background for easy identification. Proteins which can be traced back to ancestral cyanobacteria which lived 1022 mya are highlighted in yellow and orange. The phylogeny is displayed unrooted. Each node in the tree with a black circle has a posterior probability more than 0.95. Branch lengths represent amino acid substitutions with the scale bar representing an average of 1 substitution per site.

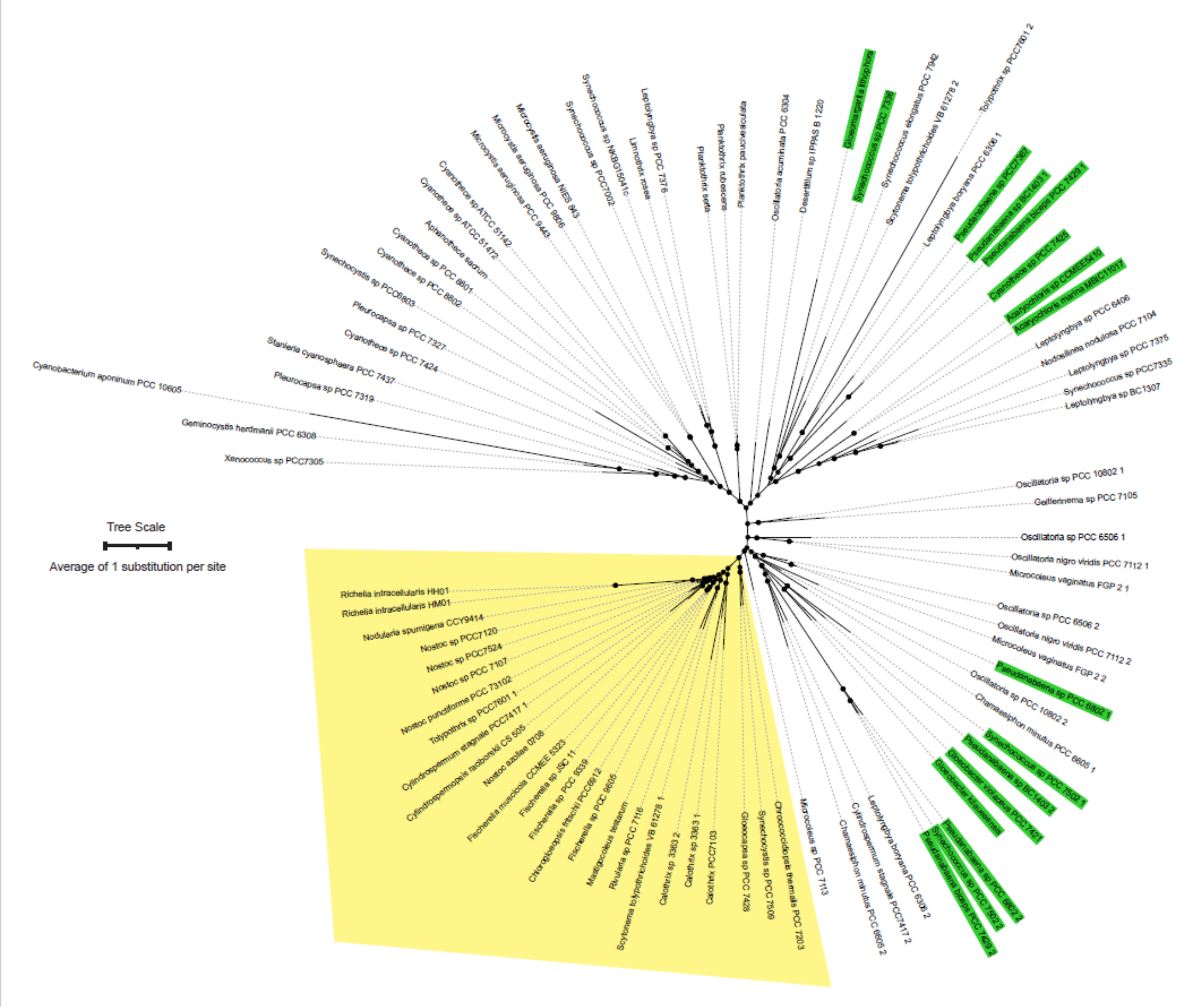


**Supplementary Figure 3. Unrooted Bayesian phylogeny of Cyanobacterial FTR1.** Protein sequences obtained using BLAST P and the reference gene from *Synechocystis* sp. PCC 6803 (Katoh et al. 2000) were aligned in MUSCLE (Edgar 2004) implemented in MegaX (Kumar et al. 2018) 396 amino acids. The alignment was truncated to 312 amino acids by removing regions of low homology. Phylogenetic reconstruction was performed in MrBayes v3.2.7a (Ronquist et al. 2012) using the mixed substitution model option with invariant sites and gamma distributed rates. Taxon labels of basal strains are labelled with a green background for easy identification. Proteins which can be traced back to ancestral cyanobacteria which lived 1822 mya are highlighted in yellow . The phylogeny is displayed unrooted. Each node in the tree with a black circle has a posterior probability more than 0.95. Branch lengths represent amino acid substitutions with the scale bar representing an average of 1 substitution per site.

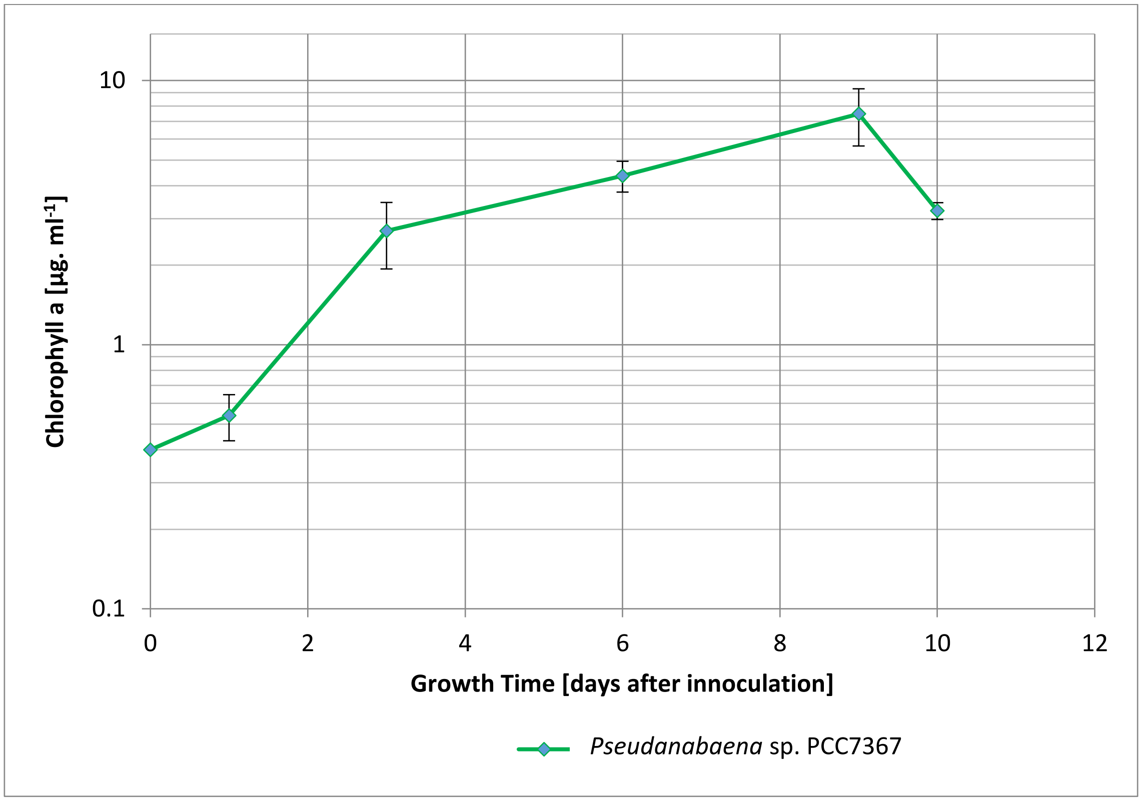


**Supplementary Figure 4: Growth of the anaerobic cultures of *Pseudanabaena* sp. PCC7367 monitored by Chl a**. The triplicate cultures were inoculated in anoxic ASN III media at 0,4 µg. $ml^{-1}$ Chl a on day 0 from exponentially growing cultures of *Pseudanabaena* sp. PCC 7367. The Chl a concentrations were averaged over the biological triplicates (n=3) with error bars indicating the standard deviation.

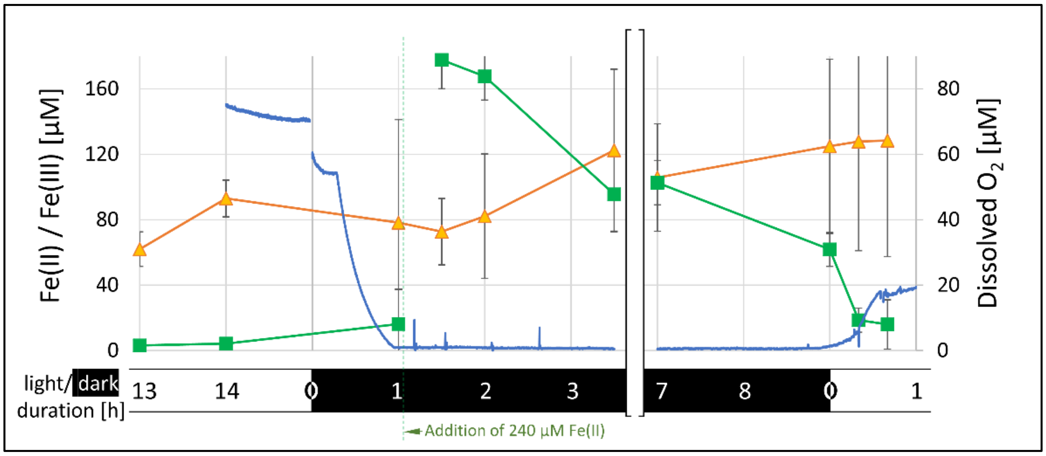


**Supplementary Figure 5:** Plot of the precipitated extracellular mean Fe(III) levels (brown line) with SD in the flask during the qPCR experiment, which was omitted from Fig. 2 for clarity. Two ml samples of the resuspended cultures were taken, centrifuged and the pellet used for Fe(III) level determination using the ferrozine assay at the time points indicated. The mean Fe(II) concentration with SD (green) was determined using the supernatant with a ferrozine assay (n = 3). The concentration of dissolved oxygen of one culture was tracked using an oxygen microsensor (blue). The point of Fe(II) addition is indicated by the green dotted line.

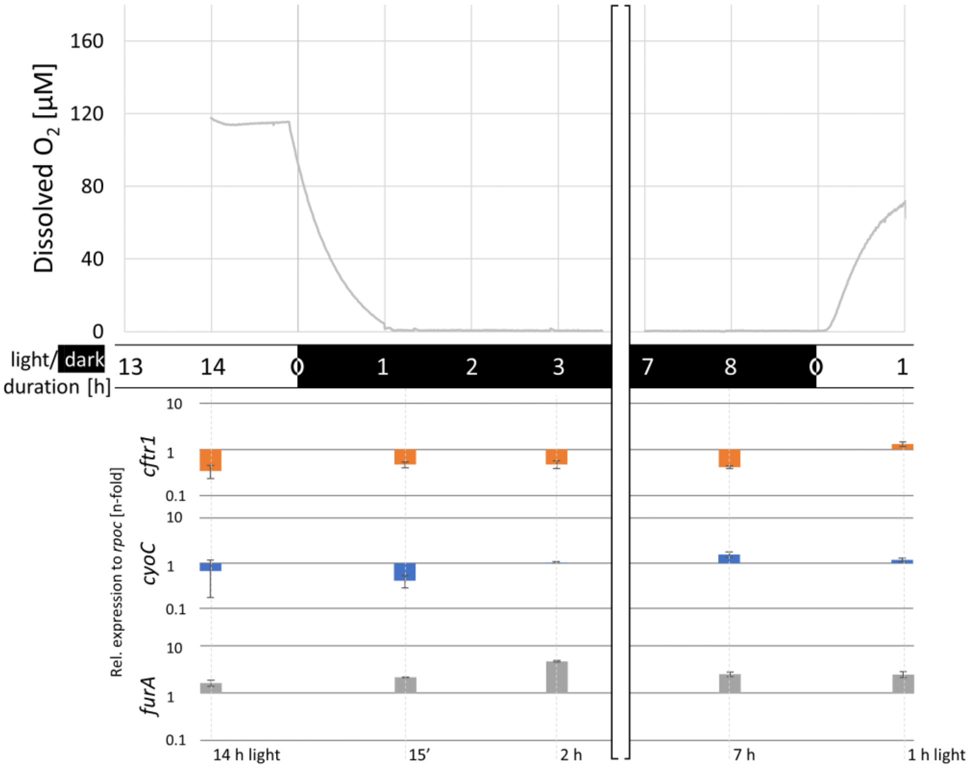


**Supplementary Figure 6: Concentrations of dissolved oxygen over the duration of the control experiment and relative qPCR gene expression data.** The concentration of dissolved oxygen in one culture (grey line), was tracked using an oxygen microsensor. Samples for RT-qPCR analysis were taken at timepoints corresponding to the experiment as indicated in Fig. 3. The change in relative expression of *cftr1, cyoC* and *furA* to the control gene, *rpoC1*, is plotted as mean fold change with SD on the y-axes for each gene (n = 3).

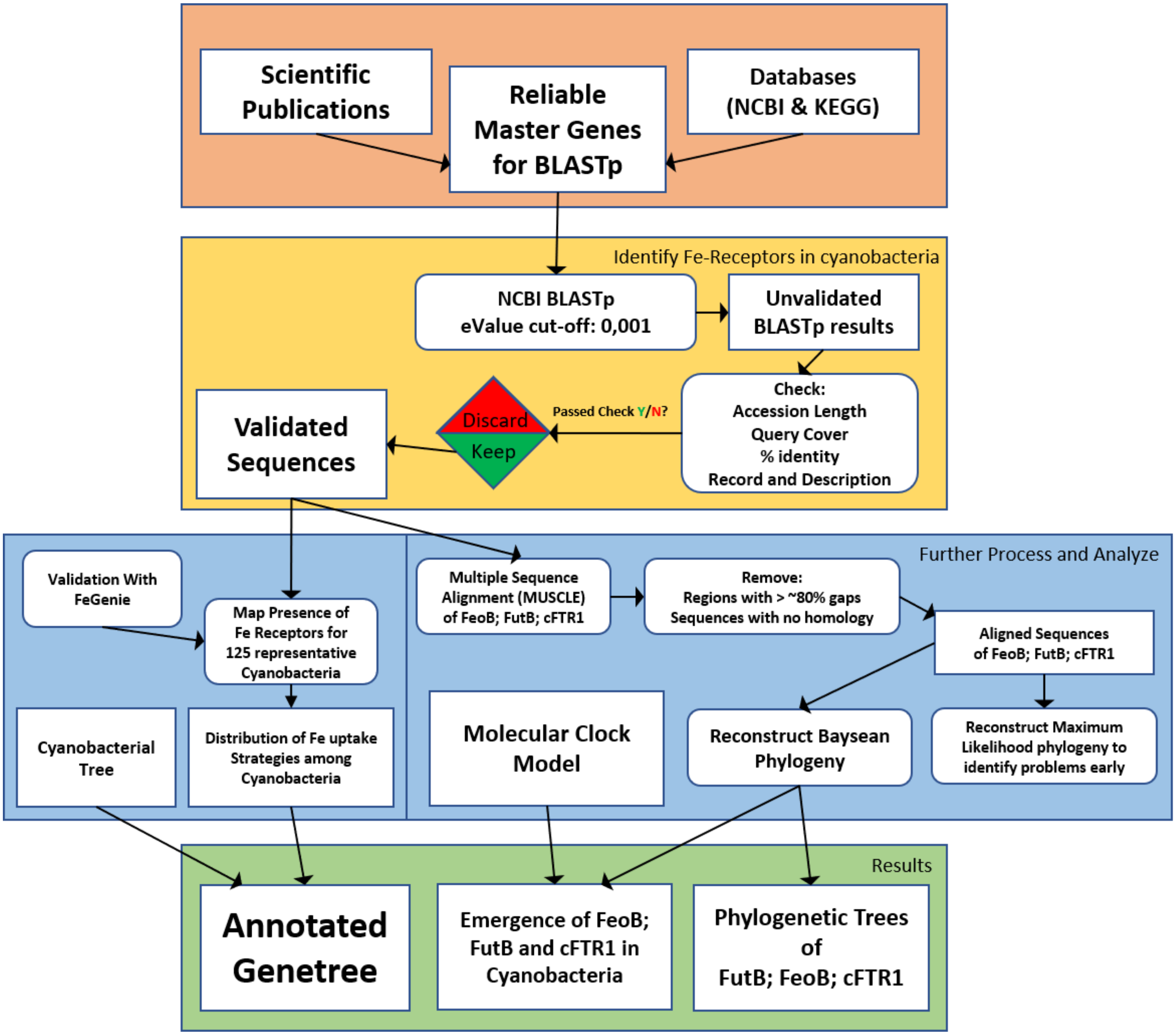


**Supplementary Figure 7: Graphical representation of the complete bioinformatics pipeline used to identify and study iron transporters in 125 Cyanobacterial genomes.** FeGenie validation was only conducted for FeoA, FeoB, FutB and cFTR1 to complement the BLASTp results. The occasions where FeGenie identified a gene undetected through BLASTp is indicated with an 'F' in Supplementary Table 2.

**Supplementary Discussion**

To predict when Fe(II) and Fe(III) transport proteins appeared in cyanobacteria, we searched for congruence between the evolutionary history of FeoB, FutB and cFTR1 and a molecular clock (Boden et al, 2021) describing the relatedness (and evolutionary histories) of different cyanobacterial strains. Therefore, monophyletic clades of a given protein which match monophyletic clades of the strains it is found within, indicate vertical inheritance from the strain's most recent common ancestor. This type of congruence between evolutionary histories of species and their iron transporters are broken when the iron transporter is transferred laterally between unrelated parental lineages. To account for this, we define 'congruence' as any group of proteins whose evolutionary history matches that of the strains they are found within. Proteins from more distant relatives can be present in this group so long as the evolutionary history of the remaining proteins mirrors a pattern of vertical inheritance in the remaining strains. Illustrated examples are presented below to demonstrate this concept:

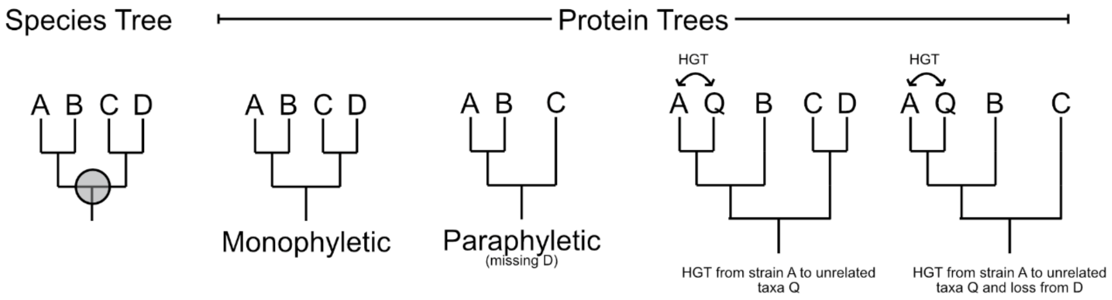


**Supplementary Figure 8:** Given the species tree, four exemplary protein trees are provided. In all scenarios, the protein (in this case an iron transporter) is assumed to have been inherited from the most recent common ancestor of strains A, B, C and D, highlighted with a grey circle in the species tree. HGT, Horizontal gene transfer
