## Supplemental Tables for "Lack of Fe(II) transporters in basal Cyanobacteria complicates iron uptake in ferruginous Archean oceans"

### Supplementary Tables

Tristan C. Enzingmüller-Bleyl[1§], Joanne Boden[2§], Achim J. Herrmann[1], Katharina W. Ebel[1], Patricia Sanchez-Baracaldo[2], Nicole Frankenberg-Dinkel[1], Michelle M. Gehringer[1*]

[1]Department of Microbiology, Technical University of Kaiserslautern, Kaiserslautern, 67663, Germany.

[2] School of Geographical Sciences, Faculty of Science, University of Bristol, Bristol, BS8 1SS, United Kingdom.

[§] These authors contributed equally to the manuscript.

*mmgehringer @yahoo.com,

Keywords: Iron uptake, Cyanobacteria, *Pseudanabaena* sp. PCC7367, Archean, evolutionary dating

### Supplementary tables

**Supplementary table 1: Iron associated transporters identified in *Pseudanabaena* sp. PCC7367.** The master genes (Column Master Gene ORF number) used in similarity searches were primarily obtained from *Synechocystis* PCC6803 (NC_000911.1) in column 'Master Gene Organism', as many publications have targeted specific iron transporters for characterisation in this species. If a similar gene was identified in *Pseudanabaena* sp. PCC7367, it is indicated with a '+' with the relative genomic open reading frame (ORF) number in the following column and the BLAST Expect value in the last column. Specifically, the following iron transporter sequences from *Synechocystis* sp. PCC6803 were used to identify homologues in *Pseudanabaena* sp. PCC7367: cyanobacterial FTR1 (Katoh *et al*. 2000), FutABC (Brandt *et al*. 2009; Katoh *et al*. 2001), FeoABC (Katoh *et al*. 2001; Kranzler *et al.* 2014) and ExbB/D TonB (Qiu *et al.* 2018; Jiang *et al.,* 2015). Genes from other Cyanobacterial species encoding the divalent metal uptake transporters classified as ZIPs (zink-iron permeases) (Morrissey & Bowler, 2012) or similar to NRAMPs (natural resistance associated macrophage proteins) (Nevo & Nelson 2006) were used as master genes for similarity searches in *Pseudanabaena* sp. PCC 7367 (NC_019701.1), namely *Chroococcidiopsis thermalis* PCC 7203 (NC_019695.1); *Synechococcus* sp. PCC7335 (NZ_DS989904.1); *Cyanobium gracile* PCC6307 (NC_019675.1), *Prochlorococcus marinus* str. AS9601 (NC_008816.1) and *Calothrix* sp. PCC7507 (NC_019682.1). For the identification of the qPCR-reference gene encoding the DNA polymerase III subunit gamma/tau (*rpoC1*) an additional master gene from *Chroococcidiopsis thermalis* PCC7203 was included, since it resulted in a more stringent expect value of 0.0 while the sequence from *Synechocystis* PCC6803 resulted in a hit with an expect value of 1e-110*.* Three amino acid sequences for subdomains of the FeoB Fe(II) transporter of *Synechocystis* PCC6803 were obtained from the KEGG genome (Kanehisa *et al*., 2016; Kanehisa &Goto, 2000) accessible at https://www.genome.jp/kegg, and these sequences are marked with 'K'. The subdomains of FeoB are: CFeoB, or the C-domain, covering amino acids 405 to 456; GFeoB, or gate domain, from amino acids 300 to 392 and 464 to 582; the NFeoB or N-domain, spanning amino acids 21 to 173 (KEGG genome). The protein sequences encoded by the master genes were used for the BLASTp (Altschul 1991; Altschul 1993; *Zhang et al.,* 2000) protein similarity search against the proteome of *Pseudanabaena* sp. PCC7367 (NC_019701.1).

| Gene | Master Gene Organism | Master Gene ORF number | *Pseudanabaena* sp. PCC7367 | ORF number | Expect Value |
|---|---|---|---|---|---|
| *cftr1* | *Synechococcus* PCC6803 | SGL_RS03385 | + | PSE7367_ RS12485 | $1E^{-71}$ |
| *fur* | *Synechococcus* PCC6803 | SGL_RS18290 | + | PSE7367_ RS06445 | $7E^{-63}$ |
| *futB* | *Synechococcus* PCC6803 | SGL_RS12585 | + | PSE7367_ RS05695 | 0 |
| | | | + (annotation in INDSEC) | Pse7367_ 1153 | 0 |
| *feoB* | *Synechocystis* sp. PCC6803 substr. PCC-P | SGL_RS05125 | - | / | / |
| *CfeoB* | *Synechocystis* PCC6803 | KEGG | - | / | / |
| *GfeoB* | *Synechocystis* PCC6803 | KEGG | - | / | / |
| *NfeoB* | *Synechocystis* PCC6803 | KEGG | - | / | / |
| *ZIP* | *Chroococcidiopsis thermalis* PCC7203 | CHRO_RS27220 | - | / | / |
| *ZIP* | *Synechococcus* sp. PCC7335 | S7335_RS13260 | - | / | / |
| *NRAMP* | *Cyanobium gracile* PCC6307 | CYAGR_RS15885 | - | / | / |
| *NRAMP* | *Prochlorococcus marinus* str. AS9601 | A9601_RS15195 | - | / | / |

| Gene | Master Gene Organism | Master Gene ORF number | *Pseudanabaena* sp. PCC7367 | ORF number | Expect Value |
|---|---|---|---|---|---|
| *NRAMP* (*mtnH*) | *Calothrix* sp. PCC7507 | CAL7507_RS21500 | - | / | / |
| *rpoC1* | *Synechocystis* PCC6803 | SGL_RS06875 | + | PSE7367_RS07505 | $1E^{-110}$ |
| *rpoC1* | *Chroococcidiopsis thermalis* PCC7203 | CHRO_RS23925 | + | PSE7367_RS07505 | 0 |
| *exbB* | *Synechocystis* PCC6803 | SGL_RS02000 | + | PSE7367_RS16055 | $1E^{-78}$ |
| | | | + | PSE7367_RS01060 | $4E^{-28}$ |
| *exbB* | *Synechocystis* PCC6803 | SGL_RS03845 | + | PSE7367_RS01060 | $8E^{-52}$ |
| | | | + | PSE7367_RS16055 | $3E^{-39}$ |
| *exbB* | *Synechocystis* PCC6803 | SGL_RS15490 | + | PSE7367_RS01060 | $3E^{-47}$ |
| | | | + | PSE7367_RS16055 | $8E^{-39}$ |

| Gene | Master Gene Organism | Master Gene ORF number | *Pseudanabaena* sp. PCC7367 | ORF number | Expect Value |
|---|---|---|---|---|---|
| *exbD* | *Synechocystis* PCC6803 | SGL_RS01995 | + | PSE7367_ RS16060 | $1E^{-34}$ |
| | | | + | PSE7367_ RS17165 | $3E^{-19}$ |
| *exbD* | *Synechocystis* PCC6803 | SGL_RS03850 | + | PSE7367_ RS17165 | $2E^{-21}$ |
| | | | + | PSE7367_ RS16060 | $3E^{-18}$ |
| *tonB* | *Synechocystis* PCC6803 | SGL_RS02005 | - | / | / |
| *tonB* | *Synechocystis* PCC6803 | SGL_RS14560 | - | / | / |
| TBDT (*fhuA*) | *Synechococcus* sp. PCC7002 | SYNPCC7002_RS15715 | + | PSE7367_ RS16045 | $2E^{-149}$ |
| | | | + | PSE7367_ RS16090 | $2E^{-137}$ |
| TBDT (*iutA*) | *Synechocystis* PCC6803 | SGL_RS01950 | + | PSE7367_ RS16090 | $2E^{-28}$ |
| | | | + | PSE7367_ RS16045 | $3E^{-26}$ |
| | *Synechocystis* PCC6803 | SGL_RS01975 | + | PSE7367_ RS16045 | 0 |

| **Gene** | **Master Gene Organism** | **Master Gene ORF number** | ***Pseudanabaena* sp. PCC7367** | **ORF number** | **Expect Value** |
|---|---|---|---|---|---|
| TBDT (*fhuE/fhuA*) | | | + | PSE7367_RS16090 | 0 |
| TBDT (*fhuA*) | *Synechocystis* PCC6803 | SGL_RS02025 | + | PSE7367_RS16045 | 0 |
| | | | + | PSE7367_RS16090 | 0 |
| FhuB | *Nostoc* sp. PCC7120 | all0387 | + | PSE7367_RS16070 | 2E-71 |
| | | | + | PSE7367_RS16065 | 8E-47 |
| FecC | *Synechocystis* PCC6803 | SGL_RS01930 | + | PSE7367_RS16065 | 1E-116 |
| | | | + | PSE7367_RS16070 | 6E-55 |
| FecD | *Synechocystis* PCC6803 | SGL_RS01935 | + | PSE7367_RS16070 | 8E-123 |
| | | | + | PSE7367_RS16065 | 2E-56 |

**Supplementary table 2: Iron transporters and related genes identified in Cyanobacteria.** The master genes used to screen for the presence of iron transporters in *Pseudanabaena* sp. PCC7367, listed in Supplementary Table 1 above, were used in protein similarity searches against 125 select Cyanobacterial proteomes. BLASTp was executed using standard algorithm settings and an e-value cut-off of 0,001 (Altschul 1991; Altschul 1993; Zhang et al. 2000). BLASTp hits were additionally validated by comparing query length, cover and identity as well as additional available information such as description of protein function and conserved domains. If a BLAST hit had greatly different query length (>~40%), different cover and identity (compare 40% cover and 10% cover) as well as a description indicating a different protein function (compare FutB iron uptake protein and ABC-Type Ammino Acid transporter) it was judged as invalid and excluded. A plus indicates that the protein is encoded, the absence is marked with a minus sign. When the gene for the protein was not identified by BLASTp, but identified with FeGenie (Garber et al. 2020) this is indicated by 'F'. In the basal clade cyanobacterium *Acaryochloris* sp. CCMEE5410 both BLASTp and FeGenie indicate that FeoB is absent, however, the KEGG genome lists a FeoB gene for this cyanobacterium, this is indicated with a 'K'. The putative ZIP protein of *Synechococcus* sp. PCC7336 was labelled with (+) as it was detected as a hypothetical protein at the cut-off value by only one of the query sequences. The first 16 rows in bold text, represent the basal clade species as presented in Figure 2B. The remaining species are listed alphabetically. The data file used to generate this table, with all cut-off values is available at https://osf.io/7x598/?view_only=715cd38c378446ba8c3f6c924f9be9f5. The percent identity to the query sequence, E-Value and accession length are listed for the best hit in the data column labelled 'Protein ID'. The 'Non-Redundant ORFs' refer to distinct sequences in a single genome assembly. Partial sequences and subdomains within a full-length sequence are not counted separately.

| Organism | cFTR1 | Fur | FutA1 | FutA2 | FutB | FutC | FeoA | FeoB | FeoC | ZIP | NRAMP | ExbBD | TonB | TBDT |
|---|---|---|---|---|---|---|---|---|---|---|---|---|---|---|
| ***Gloeobacter violaceus*** **PCC 7421** | + | + | + | + | + | + | - | - | - | + | - | + | - | + |
| ***Gloeobacter kilaueaensis*** | + | + | - | - | - | + | - | - | - | - | + | + | - | + |
| ***Synechococcus*** **sp. PCC 7336** | + | + | + | + | + | + | - | - | - | (+) | - | + | - | + |
| ***Synechococcus*** **sp. JA-3-3Ab** | - | + | + | + | + | + | - | - | - | + | - | + | - | + |
| ***Synechococcus*** **sp. JA-2-3B'a (prime a)** | - | + | + | + | + | + | - | - | - | + | - | + | - | + |
| ***Pseudanabaena*** **sp. PCC7367** | + | + | + | + | + | + | - | - | - | - | - | + | - | + |
| ***Pseudanabaena*** **sp. PCC 6802** | + | + | - | - | - | + | - | - | - | - | + | + | - | - |
| ***Synechococcus*** **sp. PCC 7502** | + | + | + | + | + | + | - | - | - | - | - | + | - | - |
| ***Pseudanabaena*** **sp. BC1403** | + | + | - | - | - | + | - | - | - | - | - | + | + | + |
| ***Pseudanabaena biceps*** **PCC 7429** | + | + | - | - | - | + | - | - | - | - | + | + | - | + |
| ***Gloeomargarita lithophora*** | + | + | + | + | + | + | - | - | - | - | - | + | - | - |

| | | | | | | | | | | | | | | |
|---|---|---|---|---|---|---|---|---|---|---|---|---|---|---|
| ***Acaryochloris marina* MBIC11017** | **+** | **+** | **+** | **+** | **+** | **+** | **-** | **+** | **-** | **+** | **-** | **+** | **-** | **+** |
| ***Acaryochloris* sp. CCMEE5410** | **+** | **+** | **+** | **+** | **+** | **+** | **-** | **+** | **-** | **+** | **-** | **+** | **-** | **+** |
| ***Cyanothece* sp. PCC 7425** | **+** | **+** | **+** | **+** | **+** | **+** | **+** | **+** | **-** | **-** | **-** | **+** | **-** | **+** |
| ***Thermosynechococcus elongatus* BP-1** | **-** | **+** | **+** | **+** | **+** | **+** | **-** | **+** | **-** | **-** | **-** | **+** | **-** | **+** |
| ***Synechococcus* sp. PCC 6312** | **-** | **+** | **+** | **+** | **+** | **+** | **+** | **+** | **-** | **-** | **+** | **+** | **-** | **+** |
| *Aphanothece sacrum* | + | + | + | + | + | + | - | + | - | - | + | + | + | - |
| *Arthrospira maxima* CS-328 | - | + | + | + | + | + | - | + | - | - | - | + | - | - |
| *Arthrospira platensis* str. Paraca | - | + | + | + | + | + | **F** | + | - | - | - | + | - | - |
| *Calothrix* PCC7103 | + | + | - | - | - | + | + | + | - | - | - | + | - | + |
| *Calothrix sp.* 336/3 | + | + | - | - | - | + | - | - | - | - | - | + | + | + |
| *Chamaesiphon minutus* PCC 6605 | + | + | - | - | - | + | - | - | - | - | + | + | - | + |
| *Chlorogloeopsis fritschii* PCC6912 | + | + | + | + | + | + | + | + | - | + | - | + | - | + |

| | | | | | | | | | | | | | | |
|---|---|---|---|---|---|---|---|---|---|---|---|---|---|---|
| *Chroococcidiopsis thermalis* PCC 7203 | + | + | + | + | + | + | - | + | - | + | + | + | - | + |
| *Crocosphaera watsonii* WH 0003 | - | + | + | + | + | + | **F** | + | - | - | - | + | + | + |
| *Crocosphaera watsonii* WH8501 | - | + | + | + | + | + | **F** | + | - | - | - | + | + | + |
| *Cyanobacterium aponinum* PCC 10605 | + | + | + | + | + | + | + | + | - | - | - | + | + | + |
| *Cyanobacterium stanieri* PCC 7202 | - | + | + | + | + | + | + | + | - | - | + | + | + | + |
| *Cyanobium gracile* PCC 6307 | - | + | + | + | + | + | **F** | + | - | - | + | + | - | - |
| *Cyanobium* sp. PCC 7001 | - | + | + | + | + | + | **F** | + | - | - | - | - | - | - |
| *Cyanothece* sp. ATCC 51142 | + | + | + | + | + | + | + | + | - | - | - | + | + | + |
| *Cyanothece* sp. ATCC 51472 | + | + | + | + | + | + | + | + | - | - | - | + | + | + |
| *Cyanothece sp*. PCC 7424 | + | + | + | + | + | + | **F** | + | - | + | - | + | + | + |
| *Cyanothece* sp. PCC 8801 | + | + | + | + | + | + | **F** | + | - | - | - | + | + | + |

| | | | | | | | | | | | | | | |
|---|---|---|---|---|---|---|---|---|---|---|---|---|---|---|
| *Cyanothece* sp. PCC 8802 | + | + | + | + | + | + | **F** | + | - | - | - | + | + | + |
| *Cylindrospermopsis raciborskii* CS-505 | + | + | - | - | - | + | - | - | - | - | - | + | - | - |
| *Cylindrospermum stagnale* PCC7417 | + | + | - | - | - | + | - | - | - | - | + | + | - | + |
| *Dactylococcopsis salina* PCC 8305 | - | + | + | + | + | + | **F** | + | - | - | + | - | - | - |
| *Desertifilum* sp. IPPAS B-1220 | + | + | + | + | + | + | **F** | + | - | + | + | + | - | + |
| *Fischerella muscicola* CCMEE 5323 | + | + | - | - | - | + | - | - | - | + | - | + | + | + |
| *Fischerella* sp. JSC-11 | + | + | - | - | - | + | + | + | - | - | + | + | + | + |
| *Fischerella* sp. PCC 9339 | + | + | - | - | - | + | + | + | - | - | + | + | + | + |
| *Fischerella* sp. PCC 9605 | + | + | - | - | - | + | + | + | - | + | - | + | + | + |
| *Geitlerinema* sp. PCC 7105 | + | + | + | + | + | + | **F** | + | - | + | - | + | - | + |
| *Geminocystis herdmanii* PCC 6308 | + | + | + | + | + | + | **-** | + | - | - | - | + | + | + |

| | | | | | | | | | | | | | | |
|---|---|---|---|---|---|---|---|---|---|---|---|---|---|---|
| *Gloeocapsa* sp. PCC 7428 | + | + | + | + | + | + | - | - | - | + | - | + | + | + |
| *Halothece* sp. PCC 7418 | - | + | + | + | + | + | **F** | + | - | - | - | + | - | + |
| *Leptolyngbya boryana* PCC 6306 | + | + | + | + | + | + | **F** | - | - | + | - | + | + | + |
| *Leptolyngbya* sp. BC1307 | + | + | + | + | + | + | - | - | - | - | + | + | - | + |
| *Leptolyngbya* sp. PCC 6406 | + | + | + | + | + | + | **F** | + | - | - | - | + | - | + |
| *Leptolyngbya* sp. PCC 7375 | + | + | + | + | + | + | + | + | - | + | - | + | + | + |
| *Leptolyngbya* sp. PCC 7376 | + | + | + | + | + | + | - | - | - | - | - | + | + | + |
| *Limnothrix rosea* | + | + | + | + | + | + | - | + | - | - | - | + | - | + |
| *Lyngbya aestuarii* BL J | - | + | + | + | + | + | + | + | - | + | - | + | - | + |
| Lyngbya majuscula 3L | - | + | + | + | + | + | + | + | - | - | - | + | - | - |
| *Lyngbya* sp. PCC 8106 | - | + | + | + | + | + | + | + | - | - | - | + | + | - |
| *Mastigocoleus testarum* | + | + | + | + | + | + | **F** | + | - | - | - | + | - | + |

| | | | | | | | | | | | | | | |
|---|---|---|---|---|---|---|---|---|---|---|---|---|---|---|
| *Microcoleus chthonoplastes* PCC 7420 | - | + | + | + | + | + | **F** | + | - | + | - | + | - | + |
| *Microcoleus* sp. PCC 7113 | + | + | + | + | + | + | - | + | - | - | - | + | + | + |
| *Microcoleus vaginatus* FGP-2 | + | + | + | + | + | + | - | - | - | - | - | + | - | + |
| *Microcystis aeruginosa* NIES-843 | + | + | + | + | + | + | **F** | + | - | - | - | + | + | - |
| *Microcystis aeruginosa* PCC 9443 | + | + | + | + | + | + | **F** | + | - | - | - | + | + | - |
| Microcystis aeruginosa PCC 9806 | + | + | + | + | + | + | **F** | + | - | - | - | + | - | - |
| *Nodosilinea nodulosa* PCC 7104 | + | + | + | + | + | + | **F** | **F** | - | + | + | + | - | + |
| *Nodularia spumigena* CCY9414 | + | + | + | + | + | + | - | - | - | - | - | + | - | + |
| *Nostoc azollae* 0708 | + | + | - | - | + | + | - | - | - | - | - | + | - | - |

| | | | | | | | | | | | | | | |
|---|---|---|---|---|---|---|---|---|---|---|---|---|---|---|
| Nostoc punctiforme PCC 73102 | + | + | - | - | - | + | - | - | - | + | + | + | - | + |
| *Nostoc* sp. PCC 7107 | + | + | - | - | + | + | - | - | - | - | - | + | - | + |
| *Nostoc* sp. PCC7120 | + | + | + | - | + | + | - | + | - | + | + | + | - | + |
| *Nostoc* sp. PCC7524 | + | + | - | - | - | + | + | + | - | + | - | + | - | + |
| *Oscillatoria acuminata* PCC 6304 | + | + | + | + | + | + | - | + | - | - | - | + | - | + |
| *Oscillatoria nigro-viridis* PCC 7112 | + | + | + | + | + | + | - | - | - | - | + | + | + | + |
| *Oscillatoria* sp. PCC 10802 | + | + | - | - | + | + | **F** | + | - | - | - | + | - | - |
| *Oscillatoria* sp. PCC 6506 | + | + | + | + | + | + | + | + | - | - | - | + | - | + |
| *Prochlorococcus marinus* str. AS9601 | - | + | + | + | + | + | - | - | - | - | + | - | - | - |
| *Prochlorococcus marinus* str. MIT 9202 | - | + | + | + | + | + | - | - | - | - | + | + | - | - |

| | | | | | | | | | | | | | | |
|---|---|---|---|---|---|---|---|---|---|---|---|---|---|---|
| *Prochlorococcus marinus* str. MIT 9211 | - | + | + | + | + | + | - | - | - | - | + | - | - | - |
| *Prochlorococcus marinus* str. MIT 9215 | - | + | + | + | + | + | - | - | - | - | + | - | - | - |
| *Prochlorococcus marinus* str. MIT 9301 | - | + | + | + | + | + | - | - | - | - | + | - | - | - |
| *Prochlorococcus marinus* str. MIT 9303 | - | + | + | + | + | + | - | - | - | - | - | - | - | - |
| *Prochlorococcus marinus* str. MIT 9312 | - | + | + | + | + | + | - | - | - | - | + | - | - | - |
| *Prochlorococcus marinus* str. MIT 9313 | - | + | + | + | + | + | - | - | - | - | - | - | - | - |
| *Prochlorococcus marinus* str. MIT 9515 | - | + | + | + | + | + | - | - | - | - | + | - | - | - |
| *Prochlorococcus marinus* str. NATL1A | - | + | + | + | + | + | - | - | - | - | + | - | - | - |

| | | | | | | | | | | | | | | |
|---|---|---|---|---|---|---|---|---|---|---|---|---|---|---|
| *Prochlorococcus marinus* str. NATL2A | - | + | + | + | + | + | - | - | - | - | + | - | - | - |
| *Prochlorococcus marinus* subsp. *marinus* str. CCMP1375 | - | + | + | + | + | + | - | - | - | - | + | - | - | - |
| *Planktothrix paucivesiculata* | + | + | + | + | + | + | - | - | - | - | - | + | - | + |
| *Planktothrix rubescens* | + | + | + | + | + | + | **F** | + | - | - | - | + | - | + |
| *Planktothrix serta* | + | + | + | + | + | + | - | + | - | - | - | + | - | + |
| *Pleurocapsa* sp. PCC 7319 | + | + | + | + | + | + | - | - | - | + | - | + | + | + |
| Pleurocapsa sp. PCC 7327 | + | + | + | + | + | + | **F** | + | - | + | - | + | + | - |
| *Prochlorothrix hollandica* PCC 9006 | - | + | + | + | + | + | - | - | - | - | - | + | - | - |
| *Richelia intracellularis* HH01 | + | + | + | - | + | + | - | - | - | - | - | + | - | - |
| *Richelia intracellularis* HM01 | + | - | - | - | + | + | - | - | - | - | - | + | - | - |

| | | | | | | | | | | | | | | |
|---|---|---|---|---|---|---|---|---|---|---|---|---|---|---|
| *Rivularia* sp. PCC 7116 | + | + | + | + | + | + | - | - | - | - | - | + | - | + |
| *Scytonema tolypothrichoides* VB-61278 | + | + | + | + | + | + | - | - | - | - | - | + | - | + |
| *Stanieria cyanosphaera* PCC 7437 | + | + | + | + | + | + | - | - | - | + | - | + | + | + |
| *Synechococcus elongatus* PCC 7942 | + | + | + | + | + | + | - | - | - | - | - | + | - | - |
| *Synechococcus* sp. 1G10 | - | + | + | + | + | + | **F** | + | - | - | - | - | - | - |
| *Synechococcus* sp. 8F6 | - | + | + | + | + | + | - | - | - | + | - | - | - | - |
| *Synechococcus* sp. BL107 | - | + | + | + | + | - | - | - | - | - | + | - | - | - |
| *Synechococcus* sp. BO 8801 | - | + | + | + | + | + | **F** | + | - | - | + | + | - | - |
| *Synechococcus* sp. CB0205 | - | + | + | + | **F** | + | - | - | - | + | - | - | - | - |
| *Synechococcus* sp. CC9311 | - | + | + | + | + | + | + | + | - | - | + | - | - | - |

| | | | | | | | | | | | | | | |
|---|---|---|---|---|---|---|---|---|---|---|---|---|---|---|
| *Synechococcus* sp. CC9605 | - | + | + | + | + | - | - | - | - | - | + | - | - | - |
| Synechococcus sp. CC9902 | - | + | + | + | + | - | - | - | - | - | + | - | - | - |
| *Synechococcus* sp. MW101C3 | - | + | + | + | + | + | + | + | - | - | + | + | - | - |
| Synechococcus sp. NKBG15041c | + | + | + | + | + | + | - | - | - | - | - | + | + | + |
| *Synechococcus* sp. PCC7002 | + | + | + | + | + | + | - | - | - | - | - | + | + | + |
| *Synechococcus* sp. PCC7335 | + | + | + | + | + | + | **F** | + | - | + | + | + | - | + |
| Synechococcus sp. RCC307 | - | + | + | + | + | - | - | - | - | - | - | + | - | - |
| *Synechococcus* sp. RS9916 | - | + | + | + | + | - | - | - | - | - | - | - | - | - |
| *Synechococcus* sp. RS9917 | - | + | + | + | + | + | **F** | + | - | - | + | + | - | - |
| *Synechococcus* sp. WH 5701 | - | + | + | + | + | + | **F** | + | - | - | + | - | - | - |
| *Synechococcus* sp. WH7803 | - | + | + | + | + | - | - | - | - | - | + | + | - | - |
| *Synechococcus* sp. WH7805 | - | + | + | + | + | - | - | - | - | - | + | - | - | - |

| | | | | | | | | | | | | | | |
|---|---|---|---|---|---|---|---|---|---|---|---|---|---|---|
| *Synechococcus* sp. WH8102 | - | + | + | + | + | + | - | - | - | - | + | - | - | - |
| *Synechococcus spongiarum* 142 | - | + | + | + | - | - | - | - | - | - | - | + | - | - |
| Synechococcus spongiarum 15L | - | + | + | + | - | - | - | - | - | - | - | + | - | - |
| *Synechococcus spongiarum* SP3 | - | + | + | + | - | + | - | - | - | - | - | + | - | - |
| *Synechocystis* sp. PCC6803 | + | + | + | + | + | + | + | + | - | - | - | + | + | + |
| *Synechocystis* sp. PCC 7509 | + | + | + | + | + | + | - | - | - | - | + | + | - | + |
| *Tolypothrix* sp. PCC7601 | + | + | - | - | - | + | **F** | + | - | - | + | + | - | + |
| Trichodesmium erythraeum IMS101 | - | + | + | + | + | + | + | + | - | - | - | + | - | - |
| *Xenococcus* sp. PCC7305 | + | + | + | + | + | + | **F** | + | - | - | - | + | + | + |

**Supplementary Table 3: Primers designed for the RT-qPCR.** The primer sequences targeting the genes under investigation: *ftr1*, a gene coding a Cyanobacterial FTR1 iron permease; *furA*, the gene coding the ferric uptake regulator and housekeeping gene, *rpoC1*, encoding the gamma subunit of the RNA polymerase in *Pseudanabaena* sp. PCC 7367. Additionally, a primer pair was designed for cytochrome c oxidase encoded by *cyoC*. The targeted gene locus, binding position from the start codon in bases, primer melting (TM) and annealing temperatures (TA) are also provided. The primer efficiencies where determined in a standard qPCR reaction using genomic DNA of *Pseudanabaena* sp. PCC7367 ranging from 10 000 000 copies to 1 copy of the targeted gene. All genes targeted were present in a single copy on the genome of *Pseudanabaena* sp. PCC7367.

| **Target Gene** | **Binding position** | | **Sequence 5' to 3'** | **Product length [bases]** | **$T_M$ [°C]** | **$T_A$ [°C]** | **Amplifi-cation efficiency** | **Detection threshold** |
|---|---|---|---|---|---|---|---|---|
| *ftr1* (Pse7367_RS12485) | cFTR1 Fw | 594 | CCGCATTATTACCCTGAGTAG | 55 | 59 | 54 | 109,6% | 10 ng |
| | cFTR1 Rv | 742 | AACGGGCATTTGTATCCTG | | | | | |
| *furA* (Pse7367_RS06445) | furA Fw | 82 | CGGGAAGTGATTCTGGATAC | 55 | 59 | 54 | 110,3% | 10 ng |
| | furA Rv | 210 | CTTCACCGATCGATAGATGG | | | | | |
| *rpoC1* (Pse7367_RS07505) | rpoC1 Fw | 1382 | CAGCGTTTAATGCTGACTTTG | 55 | 60 | 55 | 110,3% | 10 ng |
| | rpoC1 Rv | 1530 | TTGGCTAGGCGTAATAATTGG | | | | | |
| *cyoC* (Pse7367_RS00935) | cyoC Fw | 152 | CAGCCTATCTTGCCTATCG | 55 | 59 | 54 | 106,1% | 100 ng |
| | cyoC Rv | 368 | TGTCCGCCAATGAACAC | | | | | |

**Supplementary Table 4a & b: Potential outer membrane porins encoded by *Pseudanabaena* sp. PCC7367**

The permeability of the Cyanobacterial outer membrane is significantly lower than that of the heterotrophic bacterium, *E. coli* (Kowata *et al*., 2017; Qiu *et al*.,2021 for a review), requiring additional transport systems to take up nutrients, including iron. *Synechocystis* porins play a role in inorganic ion transport (Qiu *et al.* 2018), with a specific iron transporting porin recently identified (Qiu *et al*. 2021). The constitutive expression of this open reading frame under varying iron conditions suggests that it plays a role in iron homeostasis in *Synechocystis* PCC 6803 (Qiu et al. 2021). While porin encoding genes are found in numerous Cyanobacterial genomes, the iron specific transporter was not identified in *Pseudanabaena* sp. PCC7367 (Qiu *et al*., 2021). Additional potential outer membrane (OM) porins in *Pseudanabaena* PCC7367 were identified by their similarity to six putative iron porins in *Synechocystis* PCC6803 (Qiu *et al.* 2018). Screening of *Pseudanabaena* sp. PCC 7367 for the 6 putative porin genes identified in *Synechocystis* sp. PCC6803, by BLASTp (August 2021), yielded a total of 8 potential OM porins (Supplementary table 4b). The BLASTp hit with the highest percentage identity and an E-Value cut-off of 0,001 is listed in the 'Protein ID' column in Supplementary Table 4a. While all 8 OM porins identified in *Pseudanabaena* sp. PCC7367 are annotated as iron uptake porins, the putative OM porin WP_015165725.1 encoded by PSE7367_RS12540 in PCC7367 does not meet the parameters for an iron selective porin (highlighted in red in Supplementary Table 4a) as described by Qiu *et al*. (2021).

**A**

| Outer Membrane Porins | | | Pseudanabaena sp. PCC 7367 | | | | | |
|---|---|---|---|---|---|---|---|---|
| **Description** | **Query Sequence** | **ORF** | **# Hits** | **% Ident.** | **E Value** | **Length** | **Protein ID** | **ORF** |
| Putative OM porins of *Synechocystis* 6803 | WP_01087334 4.1 | SGL_RS13040 | 8 | 40.16 % | 5.00E-107 | 545 | WP_01516 5725.1 | PSE7367_R S12540 |
| | WP_01087221 3.1 | SGL_RS07085 | 8 | 35.52 % | 6.00E-94 | 545 | WP_01516 5725.1 | PSE7367_R S12540 |
| | WP_01087425 2.1 | SGL_RS17810 | 8 | 39.84 % | 4.00E-102 | 545 | WP_01516 5725.1 | PSE7367_R S12540 |
| | WP_01087402 9.1 | SGL_RS16575 | 8 | 56.85 % | 0.0 | 596 | WP_01516 5113.1 | PSE7367_R S09305 |
| | WP_01087175 8.1 | SGL_RS04720 | 8 | 40.70 % | 6.00E-138 | 545 | WP_01516 5725.1 | PSE7367_R S12540 |
| | WP_01087207 9.1 | SGL_RS06365 | 8 | 35.11 % | 1.00E-98 | 553 | WP_01516 3773.1 | PSE7367_R S02420 |

**B**

| Protein Accession | ORF | Length | Description |
|---|---|---|---|
| WP_015165725.1 | PSE7367_RS12540 | 545 | iron uptake porin |
| WP_015163773.1 | PSE7367_RS02420 | 553 | iron uptake porin |
| WP_015165113.1 | PSE7367_RS09305 | 596 | iron uptake porin |
| WP_015164617.1 | PSE7367_RS06700 | 571 | iron uptake porin |
| WP_015166173.1 | PSE7367_RS14840 | 609 | iron uptake porin |
| WP_015164709.1 | PSE7367_RS07205 | 557 | iron uptake porin |
| WP_015164616.1 | PSE7367_RS06695 | 587 | iron uptake porin |
| WP_015166634.1 | PSE7367_RS17285 | 694 | iron uptake porin |
